# A Synthetic Transcription Platform for Programmable Gene Expression in Mammalian Cells

**DOI:** 10.1101/2020.12.11.420000

**Authors:** William C.W. Chen, Leonid Gaidukov, Yong Lai, Ming-Ru Wu, Jicong Cao, Michael J. Gutbrod, Gigi C.G. Choi, Rachel P. Utomo, Ying-Chou Chen, Liliana Wroblewska, Manolis Kellis, Lin Zhang, Ron Weiss, Timothy K. Lu

## Abstract

Precise, scalable, and sustainable control of genetic and cellular activities in mammalian cells is key to developing precision therapeutics and smart biomanufacturing. We created a highly tunable, modular, versatile CRISPR-based synthetic transcription system for the programmable control of gene expression and cellular phenotypes in mammalian cells. Genetic circuits consisting of well-characterized libraries of guide RNAs, binding motifs of synthetic operators, transcriptional activators, and additional genetic regulatory elements expressed mammalian genes in a highly predictable and tunable manner. We demonstrated the programmable control of reporter genes episomally and chromosomally, with up to 25-fold more activity than seen with the EF1α promoter, in multiple cell types. We used these circuits to program the secretion of human monoclonal antibodies and to control T-cell effector function marked by interferon-γ production. Antibody titers and interferon-γ concentrations significantly correlated with synthetic promoter strengths, providing a platform for programming gene expression and cellular function in diverse applications.

## Introduction

The regulation of gene expression in complex organisms has been a central focus for characterizing disease variation [1], producing monoclonal antibodies (mAbs) [2], developing gene and cell therapies [3], and investigating other biological phenomena [4]. Synthetic biology offers powerful ways to harness artificial gene regulatory tools in mammalian systems [5–9]. For example, tumor-specific synthetic gene circuits applied to cancer immunotherapy yield significant anti-tumor responses in mice [10]. However, the use of strong constitutive promoters in gene expression platforms can increase the expression of target genes to the point of causing unwanted, dose-dependent side effects, raising safety concerns [11]. For instance, the use of the cytomegalovirus promoter (CMVp) to express a tumor-killing gene markedly increases apoptosis in normal cells and induces acute systemic toxicity *in vivo* [12]. Although promoter substitution is a simple and commonly implemented strategy for altering gene expression [13], optimizing cell type-specific or gene-specific natural promoters demands extensive effort [14]. Thus, controlling target gene expression with natural promoters has had only limited success in achieving desired biological phenotypes or therapeutic outcomes.

Another approach to regulating gene expression is to engineer transcription factors (TFs) and transcriptional activation domains (TADs) to control transcriptional activities [15]. Artificial transcription factors (aTFs), which can be rationally designed in silico, have been derived from zinc fingers (ZFs) [16], transcription activator-like effectors (TALEs) [17], and clustered regularly interspaced short palindromic repeats associated protein (CRISPR-Cas) [18, 19]. By conjugating DNA-binding proteins with various TADs, such as VP16 [20], VP64 [21], and VPR (VP64-p65-RTA) [22], researchers have demonstrated the utility of tuning promoter activity with aTFs. Particularly, CRISPR-based aTFs (crisprTFs) are simpler to customize and target to genomic loci of interest, using guide RNA (gRNA) with homology, than complex ZFs and TALEs; thus, crisprTFs are rapidly gaining popularity in biomedical research [23]. For example, several types of crisprTFs and compound activation mediators, based on deactivated CRISPR associated protein 9 (dCas), have been widely used in mammalian cells, such as dCas-VP64 [24], dCas-VPR, SunTag [25], and synergistic activation mediator (SAM) [26]. dCas-VPR, SunTag, and SAM can strongly activate genes in multiple species [27]. Our group has also developed crisprTFs to regulate gene expression driven by natural or artificial eukaryotic promoters [28].

Here, we build upon these aTF platforms by creating a comprehensive crisprTF promoter system for the programmable regulation of gene expression in mammalian cells. Our goal is to engineer a universal platform for tunable, scalable, and consistent transcriptional control in a wide variety of contexts, applicable to various cell types or target genes. Specifically, through mimicking natural mammalian promoters, we have created modular libraries of both crisprTFs and synthetic operators by: 1) altering gRNA sequences; 2) adjusting the number of gRNA binding sites (BS) in the synthetic operator; 3) incorporating additional control elements in the operator or crisprTF to augment expression; and 4) designing multiple orthogonal crisprTFs. Because it implements a multi-tier gene circuit assembly design, this system has the advantage of operating at both epigenomic and genomic levels with precise tunability, versatile modularity, and high scalability.

To demonstrate the utility of this synthetic transcription platform, we first validated the precise control of two fluorescent reporter genes and then programmed the production of recombinant human mAbs, including a functional checkpoint antibody, anti-human programmed cell death protein 1 (anti-hPD1) [29]. High-yield, stable production was achieved by using crisprTF promoters within a recombinase-mediated, multi-landing pad (multi-LP) DNA integration platform [30]. Multi-LP DNA integration in genomic safe harbor loci (e.g., the Rosa26 locus) enables predictable single-copy integration, limited transgene silencing, stable gene expression, and consistent long-term protein production [30–32]. Anti-hPD1 gene circuits expressed chromosomally with this system modulated certain anti-tumor phenotypes of human T cells. These results indicate that highly tunable, sustainable, and predictable protein expression over a wide dynamic range can be achieved with our scalable, modular synthetic transcription system.

## Materials and Methods

### Molecular cloning and genetic circuit construction

All genetic circuits in this study were constructed using a modular Gateway-Gibson assembly method [33, 34]. Briefly, gRNAs and related sequences were commercially synthesized (Integrated DNA Technologies IDT and GenScript) and then cloned into corresponding entry vectors using In-Fusion cloning (Takara Bio, San Jose, CA, USA) or a one-step Gibson reaction with the in-house master mix. Other genetic parts were cloned into corresponding entry vectors using the same approach. Multi-site LR reactions were performed between a promoter entry vector flanked by attL4 and attR1 recombination sites, a gene entry vector flanked by attL1 and attL2 recombination sites, and pZDonor_Seq(*n*)-GTW-Seq(*n+1*)_R4_R2 destination vectors containing a Gateway cassette (chloramphenicol resistance and ccdB genes flanked by attR4 and attR2 recombination sites) to generate positional expression vectors [i.e., independent transcriptional units (TUs) in different position vectors used for the subsequent Gibson assembly] [33]. Gibson reactions were performed with the Gibson Assembly Ultra Kit (SGI-DNA, La Jolla, CA, USA) at 50°C for 60 min, using equimolar concentrations (approximately 40-60 fmol per 10 μL reaction) of column-purified positional expression vectors (cleaved with I-SceI), a matching adaptor vector (cleaved with XbaI and XhoI), and a carrier vector with BxB1-attB or BxB1-GA-attB integration sites (cleaved with FseI). Gibson-assembled constructs were diluted at 1:4 and used to transform *E. coli* 10G electrocompetent cells (60080-2, Lucigen, Middleton, WI, US). Cells were selected with appropriate antibiotics on solid and liquid culture; plasmids were column-purified with QIAprep Spin Miniprep Kit (27106, Qiagen, Hilden, Germany). After verification by restriction mapping analysis and Sanger sequencing of payload TUs, correctly assembled constructs were expanded in 25 mL liquid culture with Stbl3 chemically competent cells (C737303, Invitrogen) and column-purified with QIAGEN Plasmid Plus Midi Kit (12945, Qiagen). A schematic diagram of the gene circuit construction methodology is shown in **Figure 1A**. Human elongation factor-1 alpha (EF1α) promoter and mouse CMVp were used as constitutive promoter controls. Sp-dCas9-VPR was a gift from George Church (Addgene plasmid 63798) [22]. pRRL.CMVenh.gp91.Syn.Intron.eGFP was a gift from Didier Trono (Addgene plasmid 30470), which was used for cloning the 230-bp synthetic intron sequence [35]. MS2-P65-HSF1_GFP was a gift from Feng Zhang (Addgene plasmid 61423), which was used for cloning the MS2-P65-HSF1 sequence [26]. The sequences of genetic parts and the list of plasmids are provided in **Supplemental Table 1** and **Table 2**, respectively.

**Figure 1.**
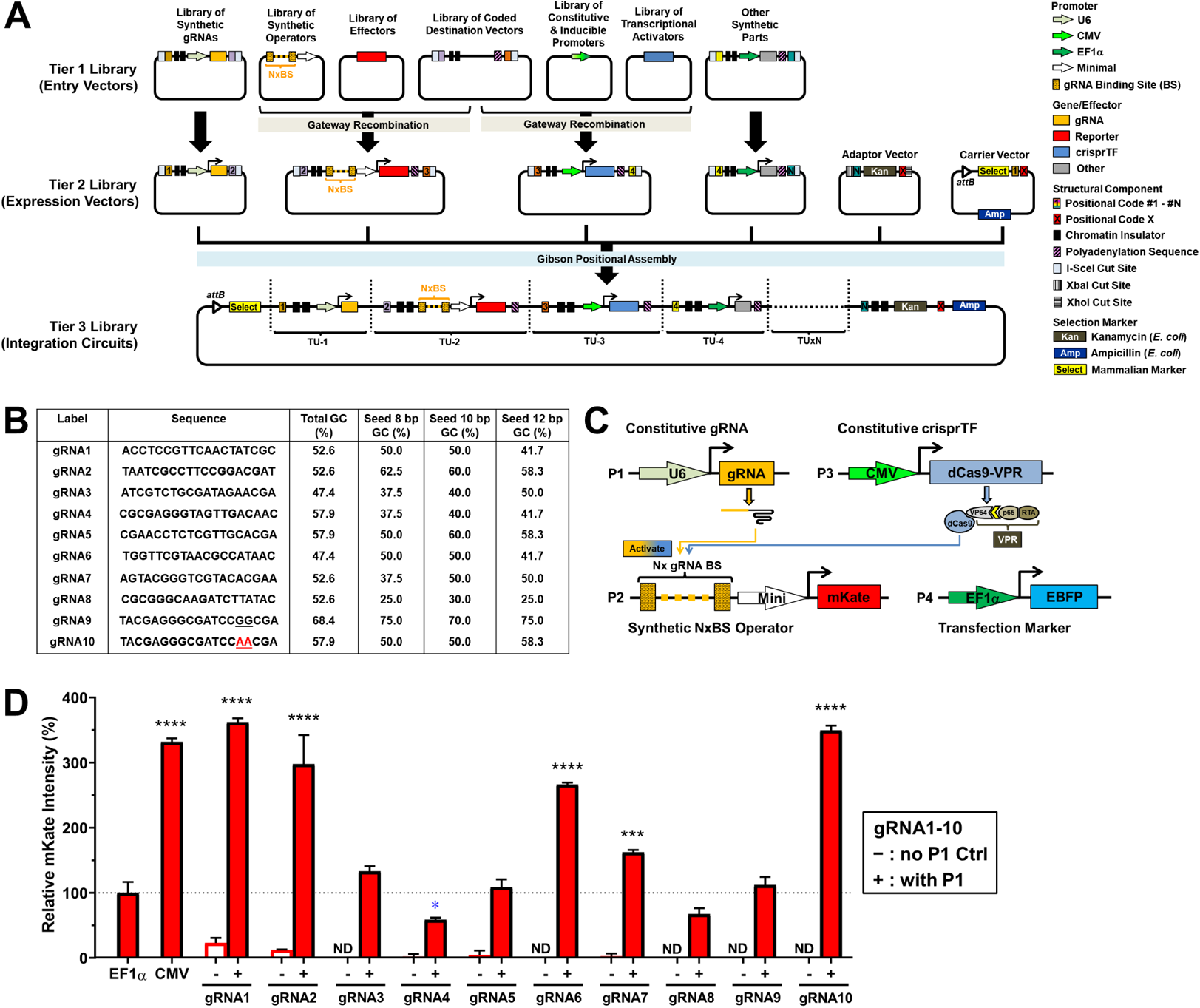
Design and development of the crisprTF promoter system. **(A)** Schematic illustration of the programmable and modular design of the crisprTF-based transcription system. To increase its programmability, this platform was modularly divided into 3 tiers of libraries constructed with the Gateway-Gibson cloning approach. The Tier 1 library was composed of entry vector modules separately encoding gRNAs, synthetic operators with gRNA binding sites (BS) upstream of a minimal promoter, effector genes, crisprTFs and their associated promoters, and other transcriptional control elements. Tier 1 library units were assembled into positional expression vectors with pre-defined orders by Gateway cloning, forming the Tier 2 library. Positional assembly by Gibson cloning was performed to connect independent transcriptional units (TUs), derived from positional expression vectors in the Tier 2 library by I-SceI restriction digestion, into complete gene circuits. The Tier 3 library comprised integration circuits enabling precision control of the target gene(s) when integrated into a landing pad (i.e., a designated chromosomal safe harbor) with BxB1 integrase-mediated, site-specific chromosomal integration. **(B)** Ten gRNAs (gRNA1-10) orthogonal to the CHO genome were screened for expression. gRNA10 was modified from gRNA9 with GG-to-AA mutations to reduce the GC content of the seed and the entire sequences. **(C)** To evaluate episomal gene expression levels, CHO-K1 cells were transiently transfected with four plasmids: plasmid #1 (P1) constitutively expressing gRNA; plasmid #2 (P2) encoding the synthetic operator with some number (x) of gRNA BS to drive mKate expression; plasmid #3 (P3) constitutively expressing a crisprTF; and plasmid #4 (P4) constitutively expressing the transfection marker (EBFP). mKate signals were assessed at 48 hours post-transfection. **(D)** For the gRNA screening, each gRNA was paired with a matching synthetic operator containing 8x gRNA BS to control mKate expression. EF1α and CMV promoters driving mKate expression served as positive controls. Experimental groups (**+**), represented by red solid bars, were transfected with four plasmids (P1-P4). Control groups (**-**), represented by red hollow bars, to detect baseline operator leakage were transfected without P1 (P2-P4). Data were normalized to the EF1α control and are presented as relative median mKate intensity (%). gRNA1 and gRNA2 operators exhibited notable leakage without gRNA, suggesting non-specific transcriptional activities that were not associated with targeted crisprTF binding. Data represent the mean ± SD (n = 3) (one-way ANOVA with multiple comparisons corrected by Dunnett test; \**p*<0.05, \*\*\**p*<0.001, \*\*\*\**p*<0.0001; ND: not detected).

### Landing pad (LP) vector construction

LP donor vectors for stable CRISPR-Cas9-mediated integration into the Chinese hamster ovary (CHO) cell genomes were constructed as previously reported [30]. Briefly, homology arm sequences for LP1-2, LP2, LP8, LP15, and LP20 sites (each arm approximately 0.5-1 kb long) were synthesized as a single gBlock (IDT) containing a PmeI restriction site between the left and right homology arms and unique BsaI cleavage sites in the 5’ and 3’ termini for Golden Gate cloning. Each gBlock was cloned into a pIDTsmart vector modified to contain compatible BsaI cloning sites. LP cassettes containing hEF1α-attP-BxB1-EYFP-P2A-Hygro (cassette 1), hEF1α-attP-BxB1-EBFP-P2A-Bla (cassette 2), or hEF1α-attP-BxB1-GA-EYFP-P2A-Hygro (cassette 3) were constructed using modular Gateway-Gibson cloning as previously described [33, 34]. LP cassettes were cloned into PmeI-linearized pIDTsmart backbones between the left and right homology arms using In-Fusion cloning (Takara Bio USA). Using this approach, we generated the following LP donor vectors: LP1-2-cassette 1, LP2-cassette 1, LP2-cassette 2, LP8-cassette 2, LP15-cassette 2, LP20-cassette 2, and LP20-cassette 3 [4, 30].

For human embryonic kidney (HEK)-293 cells, the LP donor vector was integrated into the AAVS1 locus using engineered zinc finger nucleases (ZFN) as previously described [34]. Briefly, 800 bp homology arm sequences for the AAVS1 locus, on the 5’ and 3’ ends of the ZFN cut site, were cloned into flanking position vectors. A center position vector was assembled to encode a double cHS4 core insulator, the CAG promoter, *attP* BxB1, EYFP-2A-Hygromycin, a rabbit beta-globin polyadenylation signal, and another double cHS4 core insulator. These three position vectors were verified and then assembled into the LP Shuttle Vector. A DNA fragment encoding the two DNA-binding ZF domains for the AAVS1 locus, separated by a 2A self-cleavage peptide sequence, was synthesized (GeneArt, Regensburg, Germany) and cloned into an expression vector with CAG promoter via Gateway assembly to create the ZFN-expressing vector.

### Cell culture and plasmid transfection

Adherent wild-type CHO-K1 cells (CCL-61, ATCC) and engineered CHO cells were maintained in complete HAMS-F12 (cHAMS-F12) medium (ATCC) containing 10% fetal bovine serum (FBS) (Sigma-Aldrich), 1% HyClone non-essential amino acids (GE Healthcare Life Sciences), and 1% penicillin/streptomycin (P/S) (Gibco by Life Sciences). Cells were grown in a humidified 37°C incubator with 5% CO_2_ and passaged every 2-3 days. Transfections were carried out with the Neon electroporation system (Invitrogen) with 10 µl Neon tips. Briefly, 1×10^5^ cells were suspended in R buffer and mixed well with 250 ng of individual experimental and transfection marker plasmids for 10-15 min at room temperature (RT). To ensure that all samples had the same total amount of plasmids, we supplemented the negative or positive control samples with a dummy plasmid composed of an identical expression vector backbone and a non-functional insert sequence. Cells were then electroporated with the setting of 1560 V/5 ms/10 pulses. Transfected cells were immediately transferred to a 24-well plate containing 1 mL complete culture medium without any anti-microbial reagents. Fluorescence-activated cell sorting (FACS) analysis was performed at 48 hours post-transfection. Experiments were performed with independently transfected biological triplicates.

HEK-293T cells (CRL-3216), mouse C2C12 myoblasts (CRL-1772), and rat H9c2 cardiac myoblasts (CRL-1446) were purchased from ATCC. All three cell types were thawed and expanded according to the manufacturer’s instructions and subsequently maintained in Dulbecco’s Modified Eagle medium (DMEM; 10569-044, Thermo Fisher Scientific, Waltham, MA, USA) containing 10% FBS and 1% P/S. Transfections were carried out with ViaFect transfection reagent (E4982, Promega, Madison, WI, USA). Briefly, cells were washed once with PBS, trypsinized, and re-suspended with complete culture medium at 1×10^5^ cells per 100 μL. Experimental and transfection marker plasmids (250 ng each) were added to 100 μL plain Opti-MEM medium and mixed well with ViaFect at a 1:3 ratio (total DNA by weight: ViaFect by volume), according to the manufacturer’s recommendation. After incubating at RT for 10 min, 1×10^5^ cells were added to the plasmid/ViaFect mixture and mixed well. Transfected cells were immediately transferred to a 24-well plate containing 0.8 mL complete culture medium without any antimicrobial reagents. FACS analysis was performed at 48 hours post-transfection.

The PGP1 human induced pluripotent stem cells (hiPSCs) were cultivated under sterile conditions [36]. Briefly, tissue culture plates were coated with Matrigel Matrix (354277, Corning Inc., Corning, NY, USA), following the manufacturer’s protocol, and incubated for at least 1 hour at 37°C prior to plating. hiPSCs were cultured in StemFlex medium (A3349401, Thermo Fisher Scientific), split with a five-minute treatment of StemPro Accutase (A1110501, Thermo Fisher Scientific) at 37°C every 3-4 days, and plated on Matrigel-coated plates at an appropriate density with ROCK inhibitor (1254, Tocris Bioscience, UK) at a final concentration of 10 μM for 24 hours to improve viability. hiPSCs were reverse transfected with plasmids using Lipofectamine Stem transfection reagent (STEM00001, Thermo Fisher Scientific) and Opti-MEM (31985088, Thermo Fisher Scientific) in 12-well plates in triplicate according to the manufacturer’s recommended protocol. Briefly, cells were cultured until they reached 40-50% confluence at transfection. Experimental and transfection marker plasmids (250 ng each) were added to 25 μL plain Opti-MEM I medium and mixed with 25 μL diluted Lipofectamine Stem transfection reagent (1:1 ratio), following the manufacturer’s protocol. After incubating at RT for 10 min, 50 μL DNA-lipid complexes were applied to each well. At 48 hours post-transfection with ~1×10^5^ cells per well, hiPSCs were harvested with StemPro Accutase, washed three times with PBS (100100023, Thermo Fisher Scientific), subjected to filtration with a 35-micron cell strainer, and then run on a flow cytometer for FACS analysis.

### Engineering landing pad cells

Single- and multi-LP CHO cell lines with engineered LP1-2, LP2, LP8, LP15, and LP20 loci were constructed in adherent CHO-K1 cells by homologous recombination with CRISPR/Cas9 as described [30]. Briefly, targeted integrations were performed by co-transfecting 500 ng of circular LP donor vector with 40 ng of px330-U6-chimeric_BB-CBh-hSpCas9 vector, a gift from Feng Zhang (Addgene plasmid 42230) [37], and 150 ng of U6-gRNA GeneArt DNA String (ThermoFisher). Roughly 10^5^ cells were transfected in triplicate with the Neon electroporation system with 10 µl Neon tips and seeded in 24-well plate. Cells were then transferred to a six-well plate at 3 days post-transfection and subjected to antibiotic selection with either hygromycin (600 µg/ml) or blasticidin (10 µg/ml) for two weeks, followed by clonal cell sorting with FACS. Clonal cells were verified with diagnostic PCR using locus-specific and LP-specific primers (for on-target integration) and backbone-specific primers (for off-target integration). Single-cell clones exhibiting locus-specific and backbone-free integration, as well as stable and homogenous LP expression, were expanded, banked, and subsequently used for gene circuit integration.

The single-LP HEK293 cell line was constructed with an engineered AAVS1 locus by homologous recombination with ZFN in wild-type adherent HEK293 cells as described [34]. Briefly, HEK293 cells were co-transfected with equimolar amounts of the ZFN-expressing vector and the LP Shuttle Vector, allowed to recover for 72 hours, and then selected with 200 μg/ml hygromycin for 2 weeks. Single-cell clones exhibiting locus-specific integration and stable LP expression were generated by serial dilutions of the surviving population and subsequently expanded for gene circuit integration.

### Chromosomal integration of gene circuits

For chromosomal integration of gene circuits in adherent, engineered CHO and HEK293 cells, cells were independently transfected in triplicate with 500 ng of the BxB1 integrase-expressing plasmid (pEXPR-CAG-BxB1) and 500 ng DNA of the payload plasmid, using the Neon electroporation system with the same setting as described above. Transfected cells were immediately transferred to a 24-well plate containing 1 mL complete culture medium per well without any antimicrobial reagents. At 3 days post-transfection, cells were transferred to a 6-well plate with 3 mL complete culture medium per well, followed by one of the following one-time cell selection strategies (depending on the individual payload construct designs described in the Results section and figures): 1) for single integrants in single LP (sLP), cells were selected with puromycin only (8 µg/ml for CHO and 1 µg/ml for HEK293) for gene circuits with a single selection marker, or with both puromycin (8 µg/ml) and blasticidin (10 µg/mL) for gene circuits with both selection markers; 2) for single integrants in double LP (dLP), cells were selected with either puromycin (8 µg/mL) alone or with hygromycin (250 µg/mL) and G418 (250 µg/mL) for the payload integrated in the LP2 locus, and with blasticidin (10 µg/mL) for the non-integrated, empty LP15 locus; 3) for double integrants in dLP, cells were selected with puromycin (8 μg/mL), blasticidin (10 µg/mL), hygromycin (250 µg/mL), and G418 (250 µg/mL). Following 10 days of selection, correct integration was confirmed by FACS, indicated by the complete disappearance of native LP fluorescence: either enhanced yellow fluorescent protein (EYFP), or enhanced blue fluorescent protein (EBFP), or both. Cells with chromosomally integrated gene circuits were maintained as pooled populations after selection. Cell viability and density were monitored with a Vi-CELL automated cell viability analyzer (Beckman-Coulter).

### Fluorescent imaging

All fluorescent images were taken with the EVOS FL Auto cell imaging system equipped with multiple LED light cubes, including TagBFP, GFP, YFP, RFP, and Texas Red (Thermo Fisher Scientific). Fluorescent images of the same color were taken with the same exposure settings. Scale bars were directly printed in the images.

### Flow cytometry and cell sorting

Cells were analyzed with a LSRFortessa flow cytometer, equipped with 405, 488, and 561 nm lasers (BD Biosciences). Thirty thousand events per sample were collected for analysis of median signal intensity of the transfected population, using a 488 nm laser and 530/30 nm bandpass filter for EYFP and a 405 nm laser, 450/50 filter for EBFP. Sphero rainbow calibration particles, 8 peaks (Spherotech) were used for instrument normalization and MEFL calculation. Median fluorescent intensity of the entire transfected population (i.e., all cells positive for both experimental and transfection marker signals) or the entire chromosomally integrated cell pool was measured in histograms (for single color) or dot-plots (for two or more colors). The same gating was used in all experiments (cutoff value set at 200 A.U. on the x-axis of the histogram or on both x- and y-axis of the dot-plot). Data were analyzed with FACSDiva software (BD Biosciences), FlowJo, and FCS Express 6. Flow cytometry data were largely normalized to the EF1α control and presented as relative median signal intensity (%). Some fluorescent intensity data shown as A.U. (not % of the EF1α control) represent the original readout of the fluorescent intensity from the flow cytometer without normalizing to the control group. Cell sorting was performed on a FACSAria cell sorter in the Swanson Biotechnology Center Flow Cytometry Facility at the Koch Institute (Cambridge, MA, USA). Untransfected CHO-K1 and HEK293 cells and unintegrated single- and multi-LP cells were used to set the gating. Different selection and sorting schemes were applied to target payload integration into specific LP sites, including: 1) single positive/single negative (EYFP+/EBFP− or EYFP−/EBFP+), 2) double-negative (EYFP−/EBFP−), or 3) double-negative/mKate-positive (EYFP−/EBFP−/mKate+) to select multi-LP cells with the payload(s) integrated into each designated LP. For pooled cell sorting, all sorted cells were initially seeded at 5,000-10,000 cells/cm^2^ in 24-well plates with 1 mL complete culture medium containing no antibiotics. For clonal cell sorting, single cells were initially sorted into flat-bottom 96-well plates with 100 μL complete culture medium containing no antibiotics for clonal expansion and subsequently selected and expanded in 24-well and 6-well plates and T75 flasks.

### RNA Extraction and RT-qPCR analysis

Chromosomally integrated CHO cells (3×10^6^) were collected for RNA extraction. Total RNA was extracted using the RNeasy Plus kit (74034, Qiagen) according to the manufacturer’s instructions. cDNA synthesis was performed using an iScript cDNA synthesis kit (Bio-Rad, Hercules, CA). Quantitative real-time PCR (RT-qPCR) was carried out in a LightCycler 96 System (Roche, Basel, Switzerland) using a KAPA SYBR FAST qPCR kit (KAPA Biosystems, Wilmington, MA, USA) according to the manufacturer’s instructions. The primer sequences of mKate and GAPDH for RT-qPCR are shown in **Supplemental Table 1**.

### Human antibody production and detection

A genetic circuit expressing one copy of dCas-VPR was first integrated into dLP-CHO cell lines. Cells were selected once with hygromycin (250 µg/mL) and G418 (250 µg/mL) for 10 days, followed by FACS sorting into a pool of single-positive cells using the LP-fluorescent reporters. A second, crisprTF-driven gene circuit expressing a human mAb, either JUG444 (Pfizer) or an anti-hPD1 (5C4) [10], was then integrated into the sorted cells. Doubly integrated cells were selected once by incubating them for 10 days with puromycin (8 µg/mL), blasticidin (10 µg/mL), hygromycin (250 µg/mL), and G418 (250 µg/mL), followed by FACS sorting into a pool of double-negative (EYFP−/EBFP−) cells using the LP-fluorescent reporters. All experiments were performed with 3 independently transfected biological replicates. For mAb expression analysis, 1.5×10^5^ cells were seeded in 24-well plates with 0.5 ml cHAMS-F12 medium per well without any antibiotics and maintained at 37°C for 4 days. Conditioned media were collected on day 4 for measurements. For long-term mAb expression analysis, master cultures were maintained and passaged accordingly for up to 5 weeks with measurements performed on the same day weekly. The amount of secreted JUG444 or anti-hPD1 was measured in duplicate with the Octet RED96 system using Protein A biosensors (ForteBio). Purified JUG444 or anti-hPD1 was used to generate a standard curve, from which the mAb titers were derived. Doubling time of each mAb-expressing cell population was estimated by proliferation of 5×10^4^ cells after 2 days of culture and subsequently compared with that of singly-occupied dLP-CHO cells without any mAb gene circuit.

### CHO-Tumor-T cell co-culture model and anti-tumor activity analysis

dLP-CHO cells with an integrated gene circuit expressing anti-hPD1 (5C4) were seeded into a 12-well plate in triplicate at a density of 1×10^5^ cells/well with 1 mL CHO cell culture medium. Cells were incubated at 37°C for 48 hours. Pre-activated human T cells and human ovarian cancer cells (OVCAR8) expressing a surface T-cell engager [10] were mixed at an effector-to-target (E:T) ratio of 20:1 (1×10^6^:0.5×10^5^ cells) in 1 mL T-cell culture medium (RPMI-1640 with 10% FBS, 1% P/S, 10 mM HEPES, 0.1 mM non-essential amino acids, 1 mM sodium pyruvate, and 50 μM 2-Mercaptoethanol) and then added to each well containing attached dLP-CHO cells. Co-cultures of three cell types (CHO, OVCAR8, and T cells) were incubated at 37°C. Cell-free supernatants were collected and processed at 24 hours and stored at −20℃. As an indicator of the T-cell response to tumor cells in the presence of actively secreted anti-hPD1 from dLP-CHO cells, interferon (IFN)-γ produced by T cells was quantified. IFN-γ concentrations in cell-free supernatants were determined by Human IFN-γ DuoSet enzyme-linked immunosorbent assay (ELISA; DY285; R&D systems, Minneapolis, MN, USA). EYFP−/EBFP+ dLP-CHO cells with no anti-hPD1 payload circuit were used as the control cells.

### Statistical analysis

All quantitative data are presented as mean ± standard deviation (SD). Statistical differences between groups were analyzed by Student’s *t*-test (for two groups), one-way ANOVA (for multiple groups), or two-way ANOVA (for long-term stability measurements) with 95% confidence interval. Statistical significance was set at *p*≤0.05. A Dunnett or a Tukey test was performed to correct for multiple comparisons in ANOVA post-hoc analysis. All statistical analyses, including correlation analysis and linear regression, were performed and graphed with GraphPad Prism 7.0 (GraphPad Software, La Jolla, CA, USA) statistics software.

## Results

### Construction of a programmable, modular synthetic transcription system with crisprTFs

To enable high tunability and versatility of the synthetic transcription system for a wide spectrum of applications, we adopted a 3-tiered modular library design [33]. The Tier 1 entry vector library encodes a variety of interchangeable gene regulatory parts and effector genes, including crisprTFs, gRNAs, operators, and other components (**Figure 1A**). Tier 1 parts can be assembled into the Tier 2 expression vector library for transient expression, followed by modular assembly into the Tier 3 integration gene circuit library (**Figure 1A**). We built a library of guide RNAs (gRNAs) that were orthogonal to the CHO genome and first selected eight of the scored ones for evaluation (**Figure 1B**). For each gRNA, we designed a corresponding operator containing 8x complementary gRNA BS to drive the expression of a far-red fluorescent reporter gene, mKate, and evaluated their performance episomally (**Figure 1C** and **Supplemental Figure 1**). Flow cytometry results showed a wide range of mKate expression levels among different gRNAs (**Figure 1D**). As active natural mammalian promoter sequences typically have a high GC content (57%) [38], we reasoned that the GC content of the protospacer adjacent motif (PAM)-proximal seed region (8-12 bases at the 3’ end of a gRNA and its BS sequence) plays a role in regulating gene expression. gRNAs with a GC content of around 50-60% in seed sequences appeared to express mKate at higher levels than gRNAs with a lower or higher GC content (**Figure 1B** and **1D**). Therefore, we selected a relatively weak gRNA (gRNA9) with a high GC content (**≥**70%) and introduced mutations in 2 consecutive bases within the seed sequence to alter its GC content. The resulting gRNA10 and its matching operator, with **≥**50% GC and still orthogonal to the CHO genome, yielded much higher mKate expression than its ancestor, gRNA9 (**Figure 1D**, *p***=**0.0004). The CMV control and gRNA1, 2, 6, 7, and 10 had higher expression (**Figure 1D**, all *p*≤0.0005), while only gRNA4 had lower expression, than the EF1α control (**Figure 1D**, *p*<0.05). The expression obtained with gRNA1, 2, and 10 was not significantly different from that obtained with the CMV control (all *p*>0.05). Overall, the top four gRNAs, with more than 2-fold higher expression than EF1α, had 50-60% GC in the first 8-10 bases of the seed sequences (**Supplemental Figure 2**). We selected three representative gRNAs that yielded weak (gRNA4), medium (gRNA7), and strong expression (gRNA10) to investigate further. A comparison of three crisprTFs (dCas-VP16, dCas-VP64, dCas-VPR) showed that dCas-VPR yielded a markedly higher expression level than dCas-VP16 or dCas-VP64 (**Supplemental Figure 3**, both *p*<0.0001), consistent with a previous report [27]. Thus, to enable the largest dynamic expression range possible, we selected dCas-VPR for subsequent investigation.

### Transcriptional programming in multiple mammalian cell types

To control gene expression at distinct levels, we built a library of synthetic operators containing 2x-16x gRNA BS for gRNA4, 7, and 10 (**Figure 1C**). Transient expression of mKate was assessed in CHO cells at 48 hours post-transfection (**Supplemental Figure 4**). Flow cytometry data showed dramatically different patterns of mKate expression among the three gRNA series (**Figure 2A**). Comparison of each gRNA series with expression levels observed with the EF1α promoter control showed: reporter activities for the gRNA4 series that ranged from 15% (2x BS) to 270% (16x BS), with 2x-8x BS driving notably lower, and 16x BS driving significantly higher, expression; reporter activities for the gRNA7 series that ranged from 26% (2x BS) to 760% (16x BS), with 2x-4x BS driving notably lower, and 8x-16x BS driving significantly higher, expression; and reporter activities for the gRNA10 series that ranged from 30% (2x BS) to 1107% (16x BS), with 6x-16x BS having significantly higher expression (**Figure 2A**). Significant correlations between expression levels and the number of BS were found with all three gRNA series (**Supplemental Figure 5A**, all *p*<0.05). Overall, with differences in gRNA sequences and the number of complementary gRNA BS in the operators, we achieved a wide dynamic range of approximately 74-fold change in the intensity of the reporter signals with these gRNA series in CHO cells. Gene expression was proportionate to the number of BS, indicating consistent tunability.

**Figure 2.**
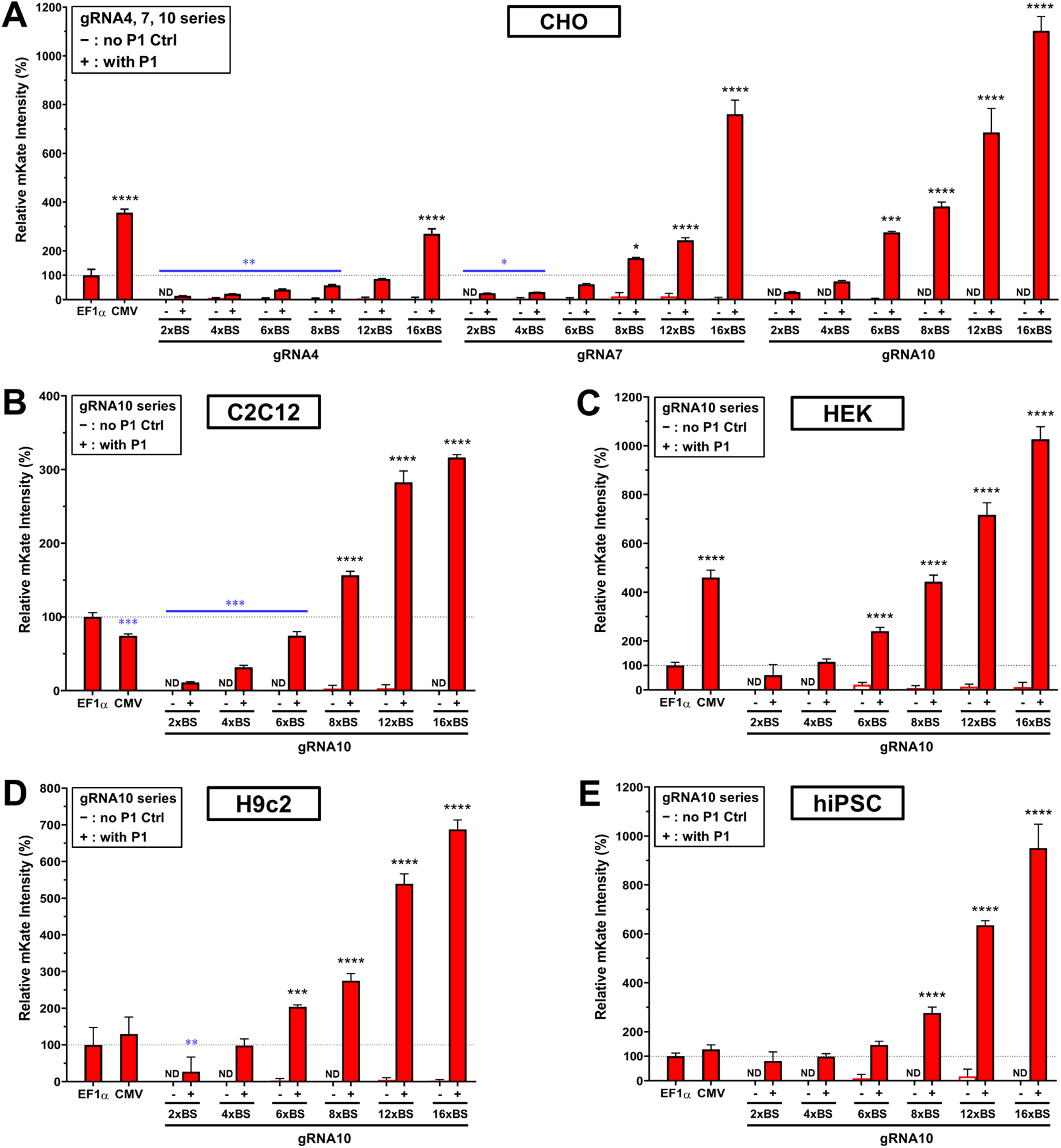
Gene expression programmed by the number of gRNA BS in synthetic operators. A library of gRNA4, 7, and 10 synthetic operators containing 2x-16x gRNA BS was built to assess the number of gRNA BS that would be effective as a programmable parameter in controlling gene expression levels. Experimental groups (**+**), represented by red solid bars, were transiently transfected with four plasmids (P1-P4 in Figure 1C) whereas gRNA-free control groups (**-**), represented by paired red hollow bars, were transfected without P1 (only P2-P4 in Figure 1C) to detect operator leakage. EF1α and CMV promoters served as positive controls. Median mKate signals relative to the EF1α control were analyzed at 48 hours post-transfection with flow cytometry. **(A)** Synthetic operators of the gRNA4, 7, and 10 series with 2x-16x BS were examined in CHO-K1 cells. **(B-E)** Synthetic operators of the gRNA10 series with 2x-16x BS were tested in mouse C2C12 myoblasts (**B**), human HEK293T cells (**C**), rat H9C2 cardiomyoblast cells (**D**), and hiPSCs (**E**). Data represent the mean ± SD (n = 3) (one-way ANOVA with multiple comparisons corrected by Dunnett test; for increased expression (marked in black): \**p*≤0.05; \*\*\**p*≤0.001; \*\*\*\**p*≤0.0001; for decreased expression (marked in blue): \**p*≤0.05; \*\**p*≤0.01; \*\*\**p*≤0.001; ND: not detected).

To translate these results into other mammalian cell types (mouse, rat, and human), we selected the gRNA10 series because it had the lowest leakage and the highest expression among the three comprehensively tested gRNAs. Interestingly, we observed dramatic differences in mKate expression with the CMV control in mouse C2C12 myoblasts (**Figure 2B**), HEK293T cells (**Figure 2C**), rat H9c2 cardiac myoblasts (**Figure 2D**), and hiPSCs (**Figure 2E**). The gRNA10 series behaved in a similar order in these cells as it did in CHO cells. In C2C12 myoblasts, mKate expression ranged from 11% (2x BS) to 316% (16x BS) of EF1α, with 2x-6x BS being notably lower and 8x-16x BS being significantly higher than that observed with EF1α (**Figure 2B**). In HEK cells, mKate expression ranged from 60% (2x BS) to 1026% (16x BS) of EF1α, with 6x-16x BS being significantly stronger than EF1α (**Figure 2C**). In rat H9c2 cells, mKate expression ranged from 27% (2x BS) to 688% (16x BS) of EF1α, with 6x-16x BS being significantly stronger than EF1α (**Figure 2D**). In hiPSC cells, mKate expression ranged from 80% (2x BS) to 950% (16x BS) of EF1α, with 8x-16x BS being significantly stronger than EF1α (**Figure 2E**). Similarly, expression levels and the number of BS markedly correlated in all four rodent and human cell types tested with the gRNA10 series (**Supplemental Figure 5B-E**, all *p*≤0.001). Collectively, with the gRNA10 series alone, we achieved up to approximately 29-fold and 17-fold changes in mKate expression in rodent and human cells, respectively.

To compare the programmable functionality of our crisprTF-based transcription platform to another orthogonal synthetic TF-based system, we constructed a Gal4-VPR-based platform containing 2x-8x UAS-BS with the same architecture (**Supplemental Figure 6A**). Although the Gal4-VPR/UAS system achieved mKate expression ranging from 11% (2x UAS-BS) to 745% (4x UAS-BS) of crisprTF 8x gRNA10-BS in CHO-K1 cells (**Supplemental Figure 6B**), we observed no correlation between gene expression levels and the number of UAS BS in the operators (**Supplemental Figure 6C**, *p*>0.05). Altogether, these results underscore the predictability and tunability of our crisprTF-based system in mammalian cells.

### Incorporation of additional genetic control elements to enhance gene expression

To extend tunability over a greater range, we incorporated the following additional genetic control elements in the crisprTF-based system: (i) a strong synthetic transcriptional activator-SAM; (ii) dual gRNA TUs to increase gRNA expression; (iii) an additional 2x nuclear localizing sequence (NLS) at the 5’ end of dCas-VPR; and (iv) a synthetic intron (SI) to enhance gene expression [39] (**Figure 3**). Transient mKate expression in CHO-K1 cells was assessed at 48 hours post-transfection. The combination of SAM, dCas-VPR, and the gRNA10 8x BS operator resulted in a 48% increase in gene expression (*p*=0.0003); however, gene expression decreased by 12.3% when SAM was combined with dCas-VPR and the 16x BS operator (*p*=0.0023), suggesting a limitation of gene activation by conjoining various TADs (**Figure 3A**). With the addition of an extra gRNA10 TU to boost the amount of gRNA, we observed modest increases, of 18.7% (*p*=0.0278) and 17.5% (*p*=0.009), with gRNA10 8x and 16x BS operators, respectively (**Figure 3B**). Thus, a single gRNA TU may be sufficient for many applications. With an additional 2x NLS incorporated at the 5’ end of dCas-VPR, mKate expression increased by 64.0% (*p*=0.0055) and 33.0% (*p*=0.0034) with gRNA10 8x and 16x BS operators, respectively, suggesting that the nuclear localization of the original dCas-VPR may be slightly insufficient to gain maximum activation of gene expression (**Figure 3C**).

**Figure 3.**
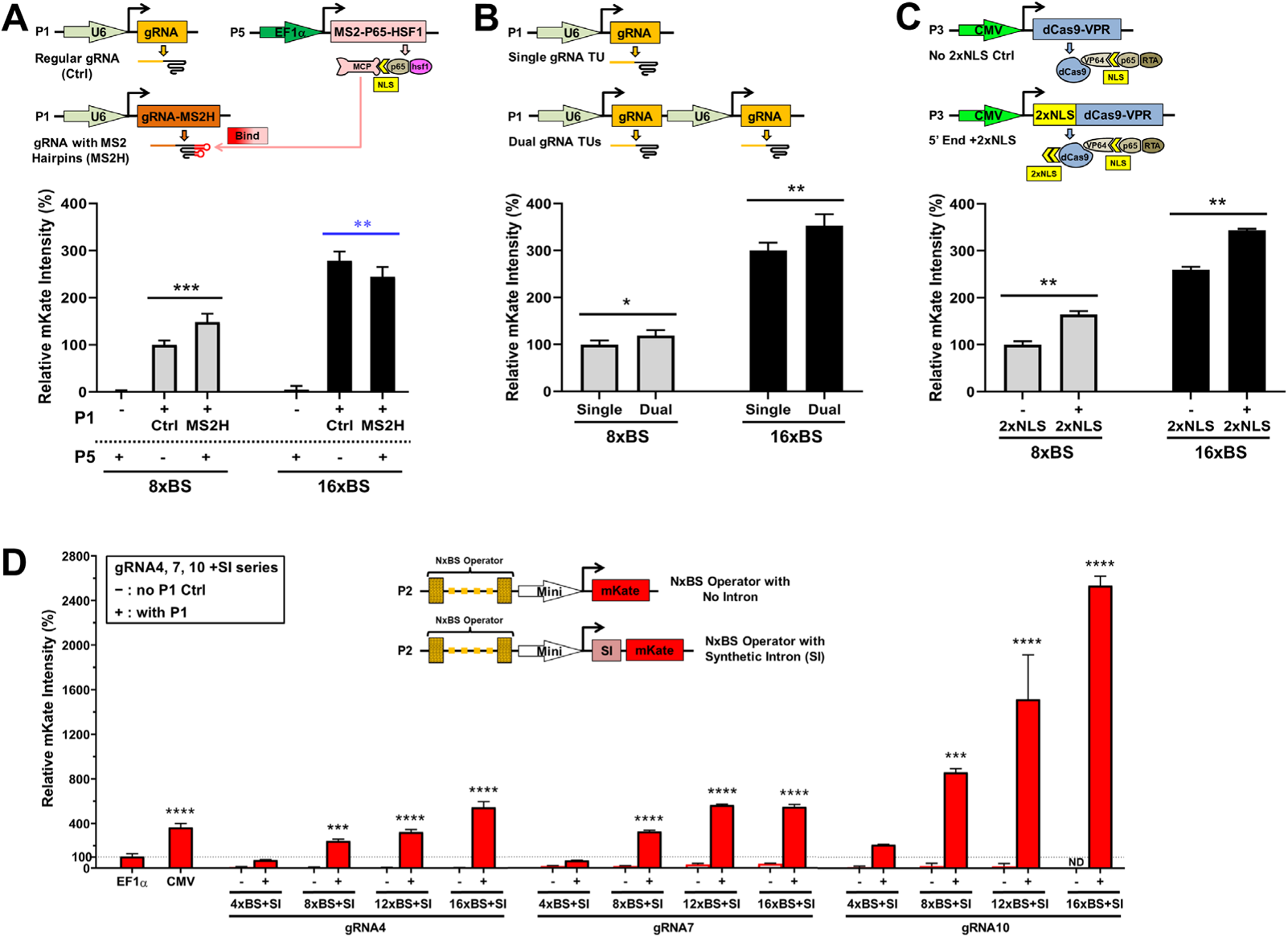
Genetic control elements added to maximize the expression level of the target gene. Schematic illustrations depicting individual experiments exhibit only plasmid constructs that were different from P1-P4 in Figure 1C. Additional genetic control elements were incorporated into the crisprTF promoters (A, B, C, and D). **(A)** Synergistic activation mediator (SAM). dCas-VPR was combined with SAM, another strong synthetic transcriptional activator. Two-tailed paired Student’s *t*-test was performed to compare P1(**+**) groups. **(B)** Dual gRNA transcriptional units (TUs). An extra gRNA10 TU was added to increase gRNA expression (two-tailed paired Student’s *t*-test). **(C)** An additional 2x nuclear localizing sequence (NLS) in the crisprTF was added at the 5’ end of dCas-VPR (two-tailed paired Student’s *t*-test). **(D)** A synthetic intron (SI) was added at the 5’ UTR of the target gene. The SI was incorporated into four synthetic operators of the gRNA4, 7, and 10 series (4x, 8x, 12x, and 16x BS), respectively. Experimental groups (**+**) were transiently transfected with four or five plasmids (P1-P4 or P1-P5), whereas negative controls (**-**) were transfected without P1. Red solid bars represent experimental groups (**+**); paired red hollow bars represent corresponding control groups without gRNA (**-**) (one-way ANOVA with multiple comparisons corrected by Dunnett test). All data represent the mean ± SD (n = 3); (for increased expression marked in black: \**p*≤0.05; \*\**p*≤0.01; \*\*\**p*≤0.001; \*\*\*\**p*≤0.0001; for decreased expression marked in blue: \*\**p*≤0.01; ND: not detected).

On the other hand, the addition of SI at the 5’ untranslated region (UTR) of the target gene led to an approximately 200-300% elevation in mKate signals with nearly all operators in all three gRNA series when compared with their original counterparts, even with the strongest operator (i.e., 16x BS of gRNA10; **Supplemental Figure 7**). The gRNA4-SI series yielded expression levels that ranged from 70% (4x BS+SI) to 528% (16x BS+SI) of those observed with the EF1α promoter, with 8x-16x BS+SI being notably higher than the EF1α control; the gRNA7-SI series ranged from 65% (4x BS+SI) to 532% (16x BS+SI) of EF1α, with 8x-16x BS+SI being significantly higher than the EF1α control; the gRNA10-SI series ranged from 205% (4x BS+SI) to 2463% (16x BS+SI) of EF1α, with 8x-16x BS+SI showing significantly higher expression than that observed with the EF1α promoter (**Figure 3D**). We did not observe any substantial increases of fluorescent signals in the absence of gRNA in any of the SI constructs (**Figure 3D**). These results indicated that SI can be fully compatible with synthetic crisprTF promoter-based gene regulation and is an efficient control element to increase gene expression. Notable correlations were found between expression levels and the number of BS with the gRNA4-SI and gRNA10-SI series (**Supplemental Figure 8**, both *p*<0.05). Taken together, by adding SI to the original gRNA operators, we expanded the achievable dynamic expression range in CHO cells to an approximately 167-fold change, offering analogue precision control of gene expression with a near-continuous spectrum (**Supplemental Figure 9**).

### Precision control of gene expression via genomic integration

We designed our crisprTF promoter platform to be fully compatible with our multi-LP DNA integration platform [30] for the stable expression of large gene circuits in engineered mammalian cells. We built gene circuits that contained insulated TUs encoding different gRNAs, gRNA operator-target gene pairs, and a crisprTF, together with an integration-enabled circuit selection marker, puromycin (**Figure 4A** and **Supplemental Figure 10**). Chromosomal integration of a single DNA copy of the gene circuit was mediated by BxB1 integrase into a single LP (sLP) locus in engineered adherent CHO (**Figure 4A**). After selection with puromycin for integration in engineered sLP-CHO cells, indicated by the disappearance of the EYFP signal, we analyzed target gene expression by flow cytometry. Both the gRNA4 and gRNA10 series exhibited tunable control of expression profiles in sLP-CHO cells, similar to that seen with their episomal counterparts (**Figure 4B**). Gene expression of the gRNA4 series ranged from 7% (2x BS) to 56% (16x BS) of that observed with the integrated EF1α control, with 2x-6x BS exhibiting markedly lower expression than with EF1α; the gRNA10 series ranged from 13% (2x BS) to 207% (16x BS) of EF1α, with 2x-4x BS also exhibiting significantly lower expression than with EF1α. Adding SI at the 5’ UTR resulted in 200-250% increases in target gene expression with representative integration circuits (**Figure 4B**, right side of the panel). With a single genomically integrated copy, only gRNA10 16x BS without SI (207%) and 8x and 16x BS with SI (223% and 521% respectively) exhibited significantly stronger expression than seen with the integrated EF1α control (**Figure 4B**). Similar to what was observed with their episomal counterparts, when both the gRNA4 and gRNA10 series were chromosomally integrated into sLP, the correlations between the number of BS in the synthetic operator and gene expression levels were highly significant (**Supplemental Figure 10A**; both *p*<0.005).

**Figure 4.**
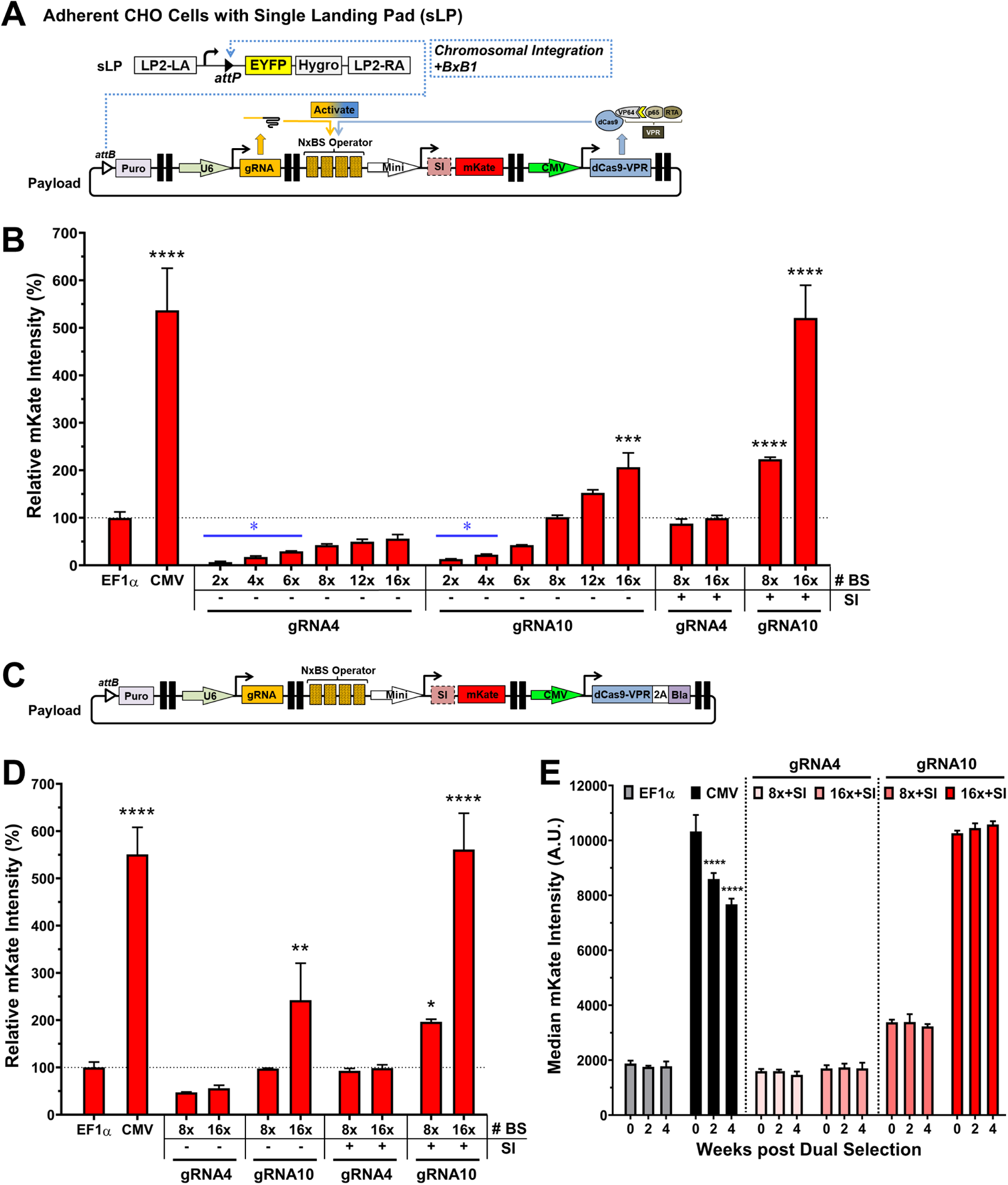
Genomic integration and long-term precision gene expression in CHO landing pad cells. **(A)** A schematic illustration of an integration gene circuit and BxB1 recombinase-mediated, site-specific integration in an engineered, adherent CHO cell line with a single landing pad (sLP). Positive integration control circuits had a central TU with an EF1α or CMV promoter driving mKate expression and two flanking dummy TUs with no gene expression in the same architecture. **(B)** The mKate signal intensities of the chromosomally integrated crisprTF promoter circuits in sLP-CHO cells relative to the integrated EF1α control circuit (one-way ANOVA with multiple comparisons corrected by Dunnett test). **(C)** A schematic illustration of an integration circuit with the addition of a 3’ flanking selection marker (blasticidin) into dCas-VPR, linked by a 2A self-cleavage peptide, to increase stability of target gene expression in long-term culture. A number of integration circuits from the gRNA4 and gRNA10 series were selected for the modification. **(D)** The mKate signal intensities of the chromosomally integrated crisprTF promoter circuits after dual selection with puromycin and blasticidin (one-way ANOVA with multiple comparisons corrected by Dunnett test). **(E)** The mKate expression levels with two of the strongest circuits from each gRNA series over the course of 4 weeks following dual selection. All four circuits examined displayed no noteworthy change in mKate signals at 2 or 4 weeks (all *p*>0.05), similar to the EF1α control (two-way ANOVA with multiple comparisons corrected by Dunnett test). All data represent the mean ± SD (n = 3) (\**p*<0.05, \*\**p*<0.01, \*\*\**p*<0.001, \*\*\*\**p*<0.0001).

To provide evidence that our crisprTF-based transcription system directly programs gene transcription, we measured the transcription levels of mKate in chromosomally integrated CHO cells. In line with mKate expression determined by flow cytometry, gene transcription of the gRNA10 series ranged from 5.7% (2x BS) to 833% (16x BS with SI) of EF1α (**Supplemental Figure 10B**). Intriguingly, with the addition of the SI, the transcription of mKate increased 6.9 folds (8x BS) and 4.3 folds (16x BS), indicating that the SI may be able to upregulate gene transcription. The correlation between the number of BS and gene transcription levels was markedly significant (**Supplemental Figure 10C**, *p*=0.0002).

To demonstrate that crisprTF circuit integration functions in human cells, we engineered a sLP in HEK293 cells and performed BxB1-mediated chromosomal integration with the gRNA10 series (**Supplemental Figure 11A**). In sLP-HEK293 cells, gene expression of the gRNA10 series ranged from 17% (2x BS) to 401% (16x BS) of EF1α (**Supplemental Figure 11B**), similar to what we recorded in sLP-CHO cells. The number of BS correlated strongly with gene expression levels in sLP-HEK293 cells integrated with the gRNA10 series (**Supplemental Figure 11C**, *p*=0.0007).

Nonetheless, by observing the expression profiles of puromycin-selected pools of crisprTF circuit integrants in sLP-CHO cells, we found that after 4 weeks of culturing, all four circuits from both the gRNA4 and gRNA10 series had notably decreased expression levels, suggesting the instability of gene expression (**Supplemental Figure 10D**, all *p*<0.05 at 4 weeks post selection). To sustain long-term expression profiles, we incorporated an additional 3’ flanking selection marker, blasticidin, into dCas-VPR, linked by a 2A self-cleavage peptide (**Figure 4C**) [40]. Flow cytometry results demonstrated gene expression levels that were similar or even slightly higher with most integration circuits immediately after one-time dual selection, compared with their unmodified counterparts (**Figure 4D**). Unlike what was observed with the CMV control (**Figure 4E**, *p*<0.0001 at 2 and 4 weeks), observations over four weeks revealed improved stability in expression levels, with no marked change for any of the four circuits examined (**Figure 4E**, all *p*>0.05 at 2 and 4 weeks). Collectively, these data suggest that a flanking selection marker co-expressed with the dCas-VPR gene may stabilize the crisprTF circuit post-integration in sLP-CHO cells.

### Modulation of human monoclonal antibody production

The controllable production of mAbs is desirable for many biomedical applications. To determine whether our transcriptional platform could be used for the precise control of antibody synthesis, we built integration gene circuits expressing a human mAb, JUG444, with the immunoglobulin kappa light chain (LC) and immunoglobulin gamma heavy chain (HC) genes separately expressed by the same gRNA10 operator as independent TUs (**Figure 5A**). We had previously observed that dCas9-VPR expression could occasionally be unstable when this construct was expressed at high levels under the control of CMVp (data not shown). To avoid the instability of dCas9-VPR expression during long-term culture, we used adherent double LP (dLP)-CHO cells, engineered with two distinct wild-type LPs: dLP1-1 and dLP1-2, to accommodate an additional copy of the dCas9-VPR gene (**Figure 5A**) [31, 41]. We first performed a targeted integration into dLP1-1 alone in dLP-CHO cells with a DNA payload encoding dCas9-VPR (**Figure 5A**). Cells were selected with three antibiotics and then subjected to single-cell FACS to isolate EYFP−/EBFP+ clones with the stably integrated dLP1-1 site and the free dLP1-2 site.

**Figure 5.**
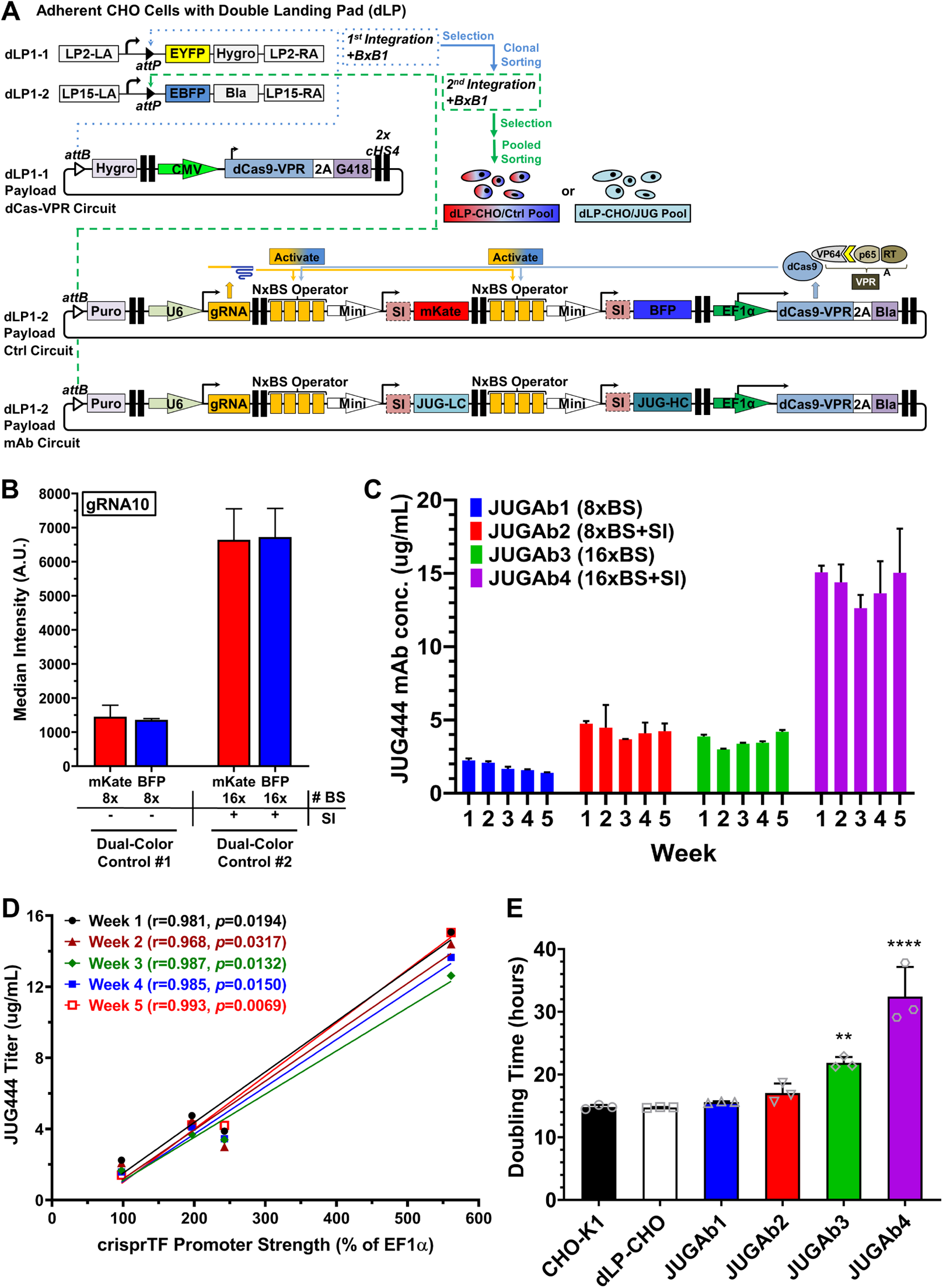
Precision control of human monoclonal antibody (mAb) production. **(A)** Schematic illustrations of the sequential, site-specific genomic integration of two payload gene circuits into the CHO cells engineered with a double landing pad (dLP) for human mAb production. First, a synthetic gene circuit encoding one copy of dCas9-VPR and 2 flanking mammalian selection markers, hygromycin (5’ end) and G418 (3’ end), was integrated into dLP1-1 site with BxB1 integrase. Selected cells with single dLP1-1 occupancy were clonally sorted based on EYFP−/EBFP+ signals. A single clone with the most consistent outputs of dCas9-VPR and EBFP was chosen for the second BxB1-mediated integration targeting the free dLP1-2 site. The second integration gene circuit contained independent TUs encoding either mKate and BFP reporter genes (control circuit) or the light chain and heavy chain of a human mAb JUG444 (mAb circuit) as well as gRNA10, dCas9-VPR, and two additional flanking selection markers: puromycin (5’ end) and blasticidin (3’ end). Integrated cells selected with four antibiotics were then pool-sorted based on EYFP−/EBFP− signals. **(B)** The mKate and TagBFP expression of the integrated payload control circuits in dLP-CHO cells with two distinct configurations (8x BS without SI and 16x BS with SI). **(C)** The mAb production of the integrated payload circuits to express the light chain and heavy chain of JUG444 with four gRNA10 operator configurations: JUGAb1 (8x BS), JUGAb2 (8x BS with SI), JUGAb3 (16x BS), and JUGAb4 (16x BS with SI). Octet mAb titer quantitation over five weeks showed stable, differential JUG444 production by all four integration circuits. **(D)** Pearson correlation analysis to determine the relationship between JUG444 mAb titers and crisprTF promoter strengths over the course of five weeks. The Pearson correlation coefficients (r) at Week 1 through Week 5 were: r=0.98 (R^2^=0.96, *p*=0.0194), r=0.97 (R^2^=0.94, *p*=0.0317), r=0.99 (R^2^=0.97, *p*=0.0132), r=0.99 (R^2^=0.97, *p*=0.015), and r=0.99 (R^2^=0.99, *p*=0.0069), respectively. **(E)** The doubling time of mAb-producing cell lines. All data represent the mean ± SD (n = 3) (one-way ANOVA with multiple comparisons corrected by Dunnett test; \*\**p*<0.01, \*\*\*\**p*<0.0001).

The functionality of clonally expanded cells was examined by integrating individual gRNA10 control circuits that co-expressed mKate and a monomeric blue fluorescent reporter, TagBFP, as well as another copy of the dCas9-VPR gene, as independent TUs into the dLP1-2 site (**Figure 5A**). Dually integrated cells were selected with four antibiotics and then FACS-sorted into pools to expand them and evaluate reporter expression (**Figure 5A**). We found that mKate and TagBFP driven by the same gRNA10 operators were simultaneously expressed at similar levels in two distinct configurations (8x BS without SI and 16x BS with SI), suggesting uniform transcriptional activation of both reporter TUs by two separate copies of the dCas-VPR gene (**Figure 5B and Supplemental Figure 12A**). Next, we replaced the control circuit in the dLP1-2 site with individual gRNA10 mAb circuits that differentially express JUG444 (**Figure 5A**). Similarly, cells with mAb circuits were selected and FACS-sorted into pools.

To examine the precision regulation and long-term stability of mAb production, we maintained cultures of sorted cell pools over five weeks and measured mAb concentration weekly, without constant antibiotic selection. We recorded differential JUG444 production in Week 1, driven by four distinct configurations: 8x or 16x BS, with or without SI (JUGAb1-4, **Figure 5C**). Stable mAb production from a single mAb gene copy was sustained throughout the experimental duration with all four configurations (**Figure 5C**). Strikingly, mAb production levels significantly correlated with the gRNA10 operator strengths documented in Figure 4D (**Figure 5D**, *p*<0.05 at all time points). The strongest operator (16x BS with SI) yielded the highest mAb level, which was comparable to the level previously observed with the strong constitutive CMVp expressing two mAb gene copies [30]. The doubling time of JUGAb3 (16x BS) and JUGAb4 (16x BS with SI) increased to 22 and 34 hours, respectively, suggesting the influence of increased antibody production on cell growth in engineered dLP-CHO cells (**Figure 5E**). Overall, these data indicated that tunable and lasting control can be applied to express human antibodies and potentially other therapeutic proteins.

### Programmable control of the human T-cell immune response against tumor cells

To explore the applications of our platform for cellular therapy, we built gRNA10 integration circuits with various configurations to program the secretion of anti-hPD1, an important immune checkpoint inhibitor widely used as a therapy in immuno-oncology [29]. The light chain and heavy chain of anti-hPD1 were separately expressed by the same gRNA10 operators as independent TUs (**Figure 6A**). Similar to the JUG444 production, a copy of dCas9-VPR was pre-integrated into the dLP1-1 site of the dLP-CHO cells to avoid its complete silencing. Each of the four circuits, designed to differentially express anti-hPD1 (8x or 16x BS, with or without SI), was then integrated into the dLP1-2 site of the dLP-CHO cells, which had been selected and clonally sorted for dLP1-1 occupancy (PD1Ab1-4, **Figure 6B**). Dually integrated cells were selected and then sorted into pools with FACS for expansion. Quantification of anti-hPD1 secretion by sorted cells showed that the PD1Ab4 group (16x BS with SI) had a significantly higher titer of anti-hPD1 than all the other groups (**Figure 6B**, all *p*<0.001) and that the PD1Ab3 group (16x BS without SI) had a notably higher titer than the PD1Ab1 group (8x BS without SI) (**Figure 6B**, *p*<0.05). A strong correlation was found between anti-hPD1 titers and crisprTF promoter strengths (**Figure 6C**, *p*<0.005).

**Figure 6.**
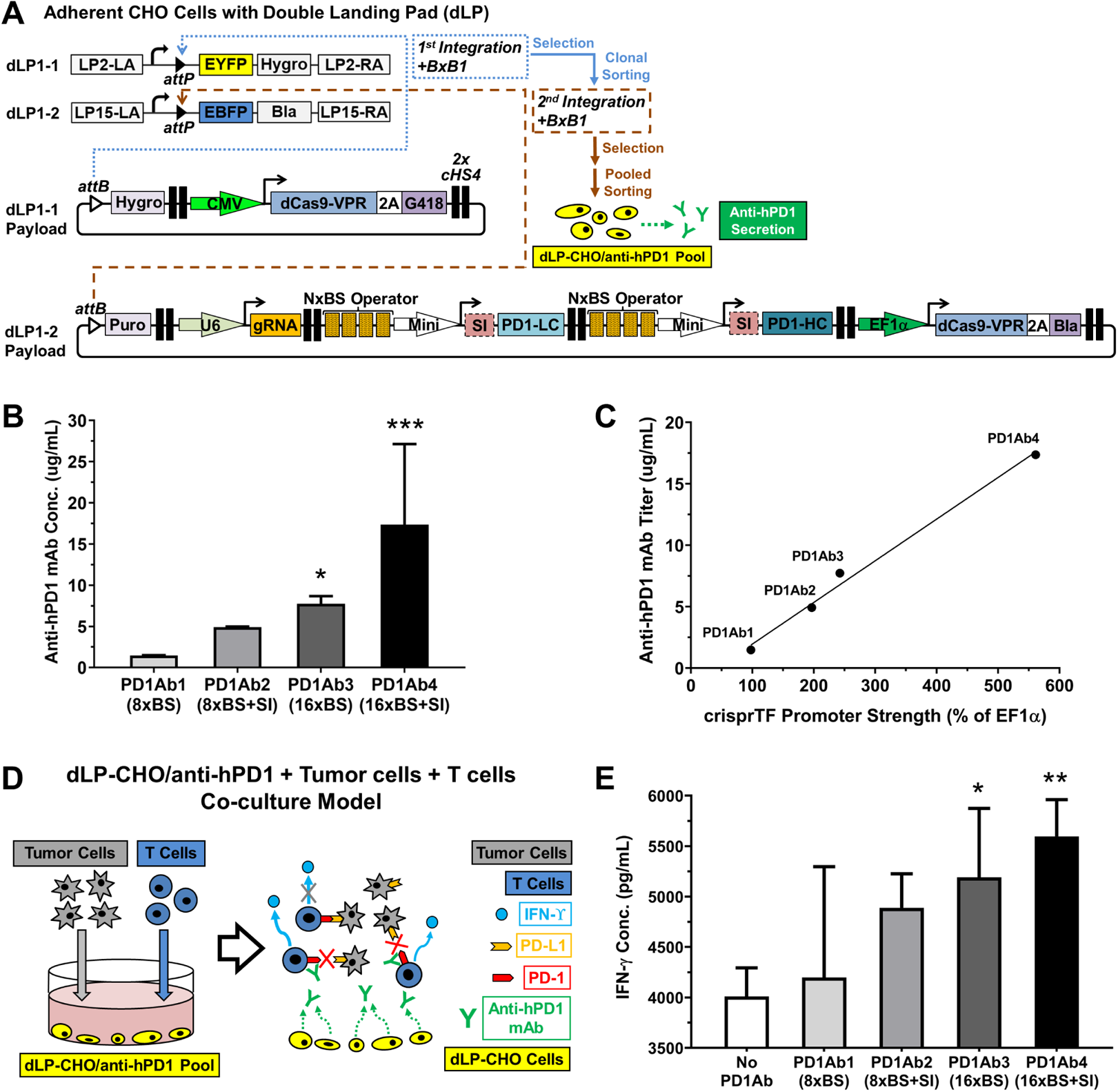
Programmable control of anti-hPD1 secretion and the human T cell anti-tumor response. **(A)** Schematic illustration of engineering the anti-hPD1-secreting dLP-CHO cells with sequential and site-specific integration of two payload gene circuits with BxB1 integrase. Clonally sorted EYFP−/EBFP+ dLP-CHO cells stably integrated with a gene circuit encoding dCas9-VPR and flanking selection markers in the dLP1-1 site were used for the second BxB1-mediated integration. The free dLP1-2 site was integrated with a gene circuit containing independent TUs that encoded: the 5’ flanking puromycin, gRNA10, the light chain and heavy chain of anti-hPD1 driven by the same gRNA10 operators, dCas9-VPR, and the 3’ flanking blasticidin linked to dCas9-VPR by a 2A self-cleavage peptide. Dually integrated cells were selected with four antibiotics and then subjected to pooled cell sorting based on EYFP−/EBFP− signals. **(B)** Octet mAb titer quantitation showed differential anti-hPD1 secretion programmed by four distinct configurations of gRNA10 operators: PD1Ab1 (8x BS), PD1Ab2 (8x BS with SI), PD1Ab3 (16x BS), and PD1Ab4 (16x BS with SI) (one-way ANOVA mixed effects analysis with multiple comparisons corrected by Tukey test). **(C)** Pearson correlation analysis revealed that anti-hPD1 titers strongly correlated with crisprTF promoter strengths (r=0.99, R^2^=0.99, *p*=0.0043). **(D)** Schematic diagram of CHO-tumor-T cell co-culture system to evaluate the functionality of anti-hPD1 and to explore the utility of our crisprTF promoter platform for cellular therapy. dLP-CHO cells engineered with one of the above four configurations for anti-hPD1 secretion were first seeded for 48 hours. The control group was seeded with EYFP−/EBFP+ dLP-CHO cells with no anti-hPD1 payload circuit. Pre-activated human T cells and human ovarian cancer cells (OVCAR8) expressing a surface T-cell engager were subsequently seeded in each well with the attached dLP-CHO cell populations. **(E)** Quantification of IFN-γ concentrations in the media at 24 hours post-co-culture by ELISA revealed tunable IFN-γ production by T cells (one-way ANOVA with multiple comparisons corrected by Dunnett test). All data represent the mean ± SD (n = 3)(\**p*<0.05, \*\**p*<0.01, \*\*\**p*<0.001).

To demonstrate the functionality of actively secreted anti-hPD1, we developed a three-way CHO-tumor-human T cell co-culture system (**Figure 6D**). We hypothesized that differentially programmed anti-hPD1 secretion would enhance T cell effector function by correspondingly blocking the interactions between tumor cells and T cells, mediated by the engagement of PD1 with programmed death ligands. Sorted, dually integrated dLP-CHO cells with individual configurations expressing anti-hPD1 were first seeded. Pre-activated human T cells and ovarian cancer cells expressing a surface-displayed T-cell engager were then added to the attached dLP-CHO cells [10]. IFN-γ production by T cells was measured at 24 hours post-co-culture as a marker of T cell activation (**Figure 6D**). T cells in the PD1Ab3 and PD1Ab4 groups produced significantly more IFN-γ than the control group (*p*<0.05 and *p*<0.01 respectively), corresponding to the higher titers of anti-hPD1 secretion (**Figure 6E**). IFN-γ production also notably correlated with anti-hPD1 titers in dLP-CHO cell cultures prior to the start of co-culturing (**Supplemental Figure 12B**, *p*<0.05). Thus, this transcription system can be used to produce functional proteins of clinical interest and perturb cellular phenotypes in a highly regulated and precise manner.

## Discussion

We have built and characterized a crisprTF promoter system for the programmable regulation of gene expression in mammalian cells. This system functions consistently across multiple mammalian cell types and is, therefore, sufficiently versatile to be used in a variety of cellular models for biomedicine and biomanufacturing.

To demonstrate the generalizability of the system’s gene regulatory activity, we used both episomal vectors in transiently transfected cells and site-specific, stably integrated chromosomal constructs in LP-engineered cells. We modulated three key parameters: 1) the gRNA sequence; 2) the number of gRNA BS repeats in the operator; and 3) the CRISPR-based transcriptional activator (different crisprTFs, different selection markers in the crisprTF, or an additional copy of the crisprTF in LP-engineered cells). We also incorporated genetic elements to enhance gene expression, including SAM, an extra gRNA TU, extra NLS, and SI in the operator. Systematic characterization of these constructs resulted in >1,000-fold range of gene expression levels. The strongest of our synthetic promoters, composed of 16x gRNA BS repeats, was significantly stronger than CMVp, one of the strongest constitutive promoters currently used for mammalian applications [13]. Recent systematic investigation of mismatched single gRNAs (sgRNAs) has revealed sgRNA-DNA interaction rules controlling endogenous gene expression and correlated phenotype [42]. In agreement with previous findings, our introduction of a 2-bp mutation that moderately altered the GC ratio in the PAM-proximal seed region of gRNA10 and its BS markedly changed its gene expression profiles, suggesting the importance of the gRNA seed sequence in controlling our crisprTF promoter-driven transcriptional activities. We anticipate that additional large-scale experimental screening, combined with computational modeling or machine learning, could be used to program gene expression even more precisely.

Various mechanisms of controlling gene expression at the transcriptional level in eukaryotes have been identified [43, 44]. Core promoters and neighboring genetic elements play instrumental roles in consolidating complex cascades of signaling events involved in transcription, impacting gene expression [43]. The complexity of combinatorial interactions among constituent TF regulatory elements hinders the creation of synthetic mammalian promoters using natural motifs. Although this issue has been gradually resolved *in silico*, it remains challenging to overcome the context- or species-dependency for the *de novo* design of mammalian promoters [44]. Positive transcriptional elements have been found between −350 and −40 bp relative to the transcription start sites (TSS) in many of the tested human promoters, whereas negative regulatory elements are more likely to be located −350 to −1000 bp upstream of the TSS [38]. Therefore, we positioned our synthetic operators at up to roughly −400 bp upstream of the TSS. Our data are consistent with previous reports suggesting that longer transcriptional bursts and thus higher expression levels might be achieved with synthetic transcriptional activators by arraying multiple BS upstream of a given promoter [28, 45]. The results of our episomal tests with all three gRNA series established strong linear correlations between gene expression levels and the number of BS (up to 16x) in the operators in multiple mammalian species and cell types (**Supplemental Figures 5** and **8**). We found similar significant correlations with two gRNA series when they were genomically integrated in LP-engineered cells (**Supplemental Figure 10A** and **11C**).

Besides tuning transcriptional activities by altering gRNA sequences and operator strengths, we discovered several compatible genetic control elements that synergistically augment gene expression. When combined with dCas-VPR, SAM moderately increased expression at medium, but not high, levels episomally. We examined the compatibility of SAM with dCas-VPR when genomically integrated, by assembling large gRNA circuits that contained independent TUs encoding both systems (**Supplemental Figure 13A**). Surprisingly, the VPR-SAM hybrid system at medium expression level generated no further increase in gene expression; in fact, at high expression level, VPR-SAM actually decreased expression (**Supplemental Figure 13B**). This seemingly paradoxical result suggested possible competition for transcriptional resources among components within the hybrid system, for example, overlapping TADs encoded in SAM and dCas-VPR genes, including VP64 and p65 [46, 47]. It remains to be tested whether alternative CRISPR-based gene activation strategies, such as the CRISPR-assisted *trans* enhancer, would be compatible for combinatorial use with our system for auxiliary augmentation of transcription [48].

On the other hand, introns, which enhance gene expression in mammalian cells when applied as a positive regulatory element [49], have been proposed to control post-transcriptional RNA processing or transport [39]. Here, the incorporation of an SI immediately downstream of the operator in the 5’ UTR of the target gene [50] increased gene expression at least 200% with nearly all tested constructs, whether episomal or chromosomal, suggesting a high degree of compatibility between the transcriptional control by the crisprTF promoters and the intron-mediated enhancement by SI [51]. RT-qPCR data also suggest that an SI in the 5’ UTR directly contributes to gene transcription in our crisprTF promoter system, consistent with gene expression enhancement by 5’ UTR sequences [52]. However, overly strong target gene expression may impact cell proliferation (**Figure 5E**). Thus, it is essential to balance target gene expression with cellular homeostasis when employing strong crisprTF promoters. Moreover, the large size of the crisprTF promoter constructs, especially with additional genetic regulatory elements, limits their use with LP-engineered cells for long-term expression. Broader applications in biomedicine may require minimizing construct size.

CHO cells, approved by regulatory agencies, are widely used by the pharmaceutical industry for the biomanufacturing of recombinant therapeutic proteins with human-like glycosylation profiles [53, 54]. CMVp and Simian virus 40 early promoter are frequently used in CHO cells to produce high titers of therapeutic proteins [55]. Yet, protein-producing cell lines often exhibit a wide range of gene expression profiles, even at the clonal level, and the mechanisms that drive viral promoters in eukaryotic cells are ill-defined [56, 57]. Natural promoters have often evolved synchronously with functionalities that depend on specific genetic contexts, so promoter activity may not transfer across various genes, conditions, or species [43, 57]. The growing number of biotherapeutic proteins in development has created an increasing demand for efficient long-term protein expression systems for biomanufacturing, ideally with built-in programmable control for improved consistency and predictable yields.

Coupled with the LP technology, we demonstrated the long-term stability of precisely tuned human mAb production with the crisprTF promoter system. Antibody yields obtained with the strongest crisprTF circuit expressing a single mAb LC/HC cassette were similar to those obtained with two or six mAb cassettes driven by CMV or EF1α promoters, respectively, in LP-engineered CHO cells [30]. The high production efficiency of our strong crisprTF circuits, along with their built-in modularity and scalability, suggests that they can be adapted to achieve industrially relevant mAb titers. Our platform may also be applied, more generally, to CHO strains optimized for industrial bioreactor conditions to: 1) increase target protein production in a predictable and controlled manner; 2) produce multiple target proteins simultaneously with specific ratios; and 3) strike a balance between cell behaviors and protein production in long-term cultures.

To exogenously modulate target gene expression by controlling gRNA transcription levels, we incorporated two types of small molecule-inducible switches into the RNA polymerase III (Pol III) promoter (**Supplemental Figure 14A**). The results in the episomal context indicate that gRNA transcription could be tuned by adjusting the concentrations of the inducers Dox and IPTG for Tet and Lac operons, respectively (**Supplemental Figure 14B** and **C**), following gene expression kinetics at different time points (**Supplemental Figure 14D**). Thus, an additional layer of target gene tunability could be achieved in our crisprTF promoter system by using inducible Pol III promoters.

In conclusion, we have built a tunable, scalable, and sustainable gene expression system of crisprTF-based regulatory elements. This platform should enable programmable, multiplexed gene modulation for broad applications, such as mammalian synthetic biology, biomanufacturing, and precision medicine.

## Acknowledgements

The authors gratefully thank Nevin M. Summers and Dr. Louane Hann for their administrative support, Kristjan E. Kaseniit for his contribution in gRNA design, Dr. Kevin J. Lebo for his expert input in the system design, Dr. Samuel D. Perli for genetic materials, Dr. Lei Wang for providing the hiPSC cell line, and Dr. Wen Allen Tseng for genetic materials and his contribution to construct design. We greatly thank Selamawit Mamo, Kalpana Jagtap, Na Li, Cong Liu, and Jonathan L. Lyles for their technical support with CHO cell cultures and assays, Mailing Ding and Christina Harrison for their technical support with molecular cloning and plasmid preparation, and Karen Pepper for editing the manuscript. We thank Dr. G. Felsenfeld (National Institute of Diabetes and Digestive and Kidney Diseases, USA) for the HS4 insulator sequences.

## Data availability

The authors declare that all relevant data supporting the findings of this study are available within the paper and its supplementary Information. Biological materials generated in this study are available on Addgene or from the corresponding author upon request.

## Supplemental Figures and Figure Legends

**Supplemental Figure 1.**
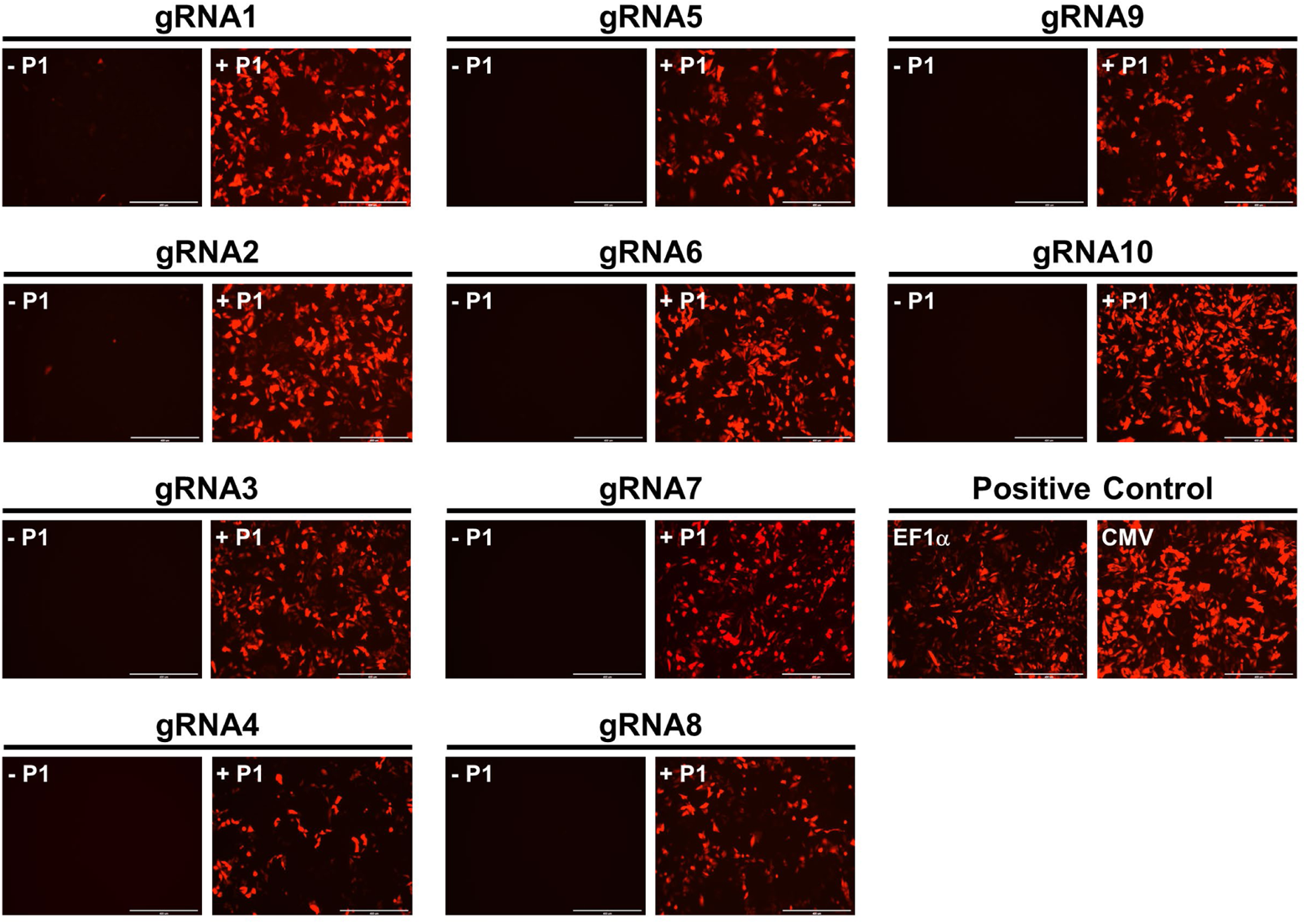
Comparison of episomal gene expression levels with 10 distinct gRNA sequences. Each gRNA was paired with a corresponding synthetic operator containing 8x gRNA BS to control mKate transcription. CHO-K1 cells were transiently transfected as illustrated in **Figure 1C**, with gRNA constitutively expressed by the U6 promoter from plasmid #1 (P1). Experimental groups were transfected with all 4 plasmids, including P1 (**+ P1**); negative control groups (no gRNA) were transfected without P1 (**−P1**). Plasmids with mKate expression driven by constitutive promoters (EF1α or CMV), transfected at the same concentration, served as positive controls. Positive and negative control groups were also supplemented with a dummy plasmid to ensure that all groups had the same total amount of transfected plasmids. Representative fluorescent images revealed a wide range of mKate expression among the 10 different gRNAs at 48 hours post-transfection. Only gRNA1 and gRNA2 exhibited slight leakage of mKate expression without the presence of P1.

**Supplemental Figure 2.**
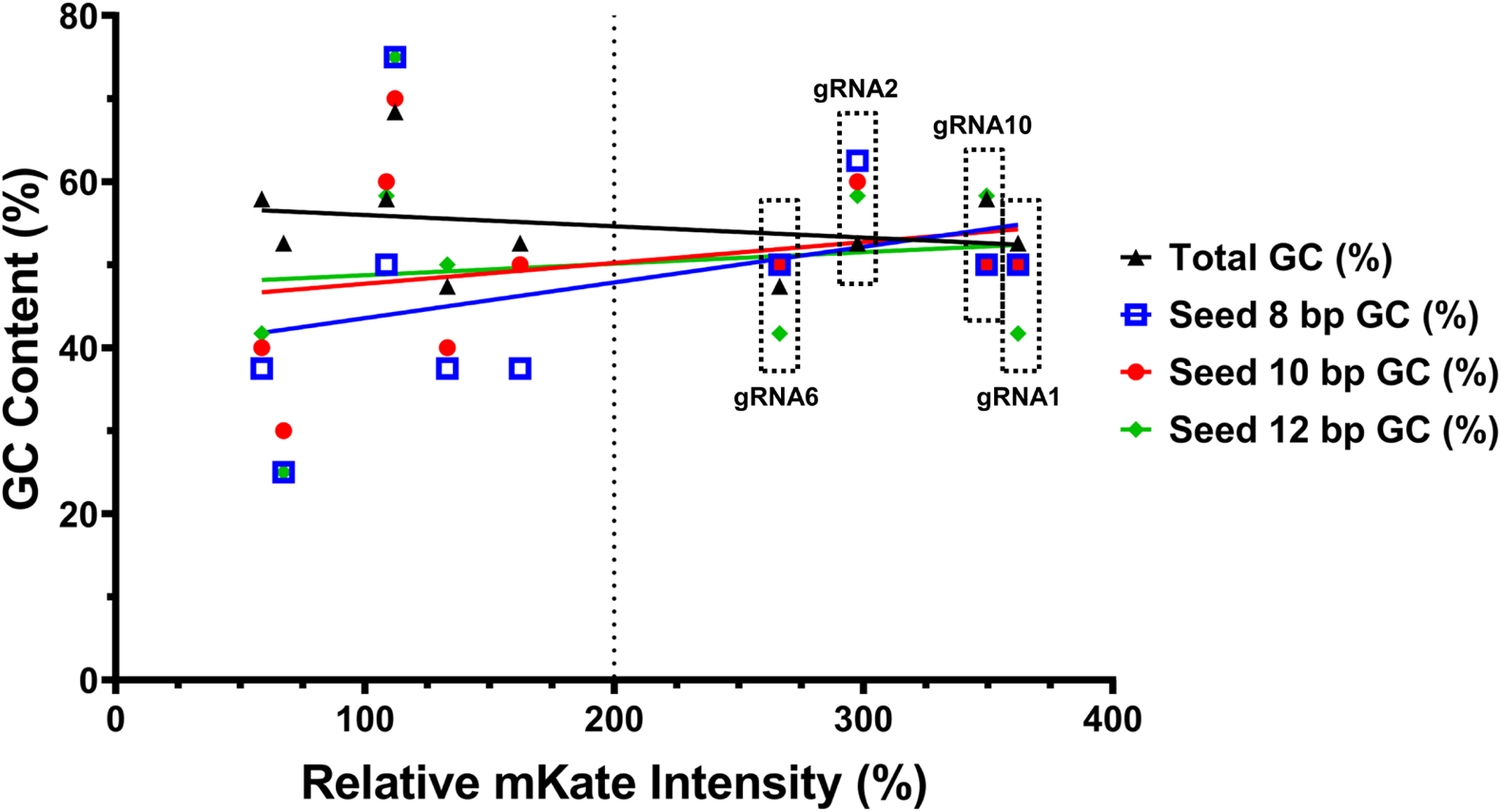
Analysis of the correlation between GC content in gRNA seed sequences and gene expression levels. We analyzed the relationship between GC content in gRNA seed 8, 10, and 12 bp sequences and corresponding mKate expression levels among gRNA1-10. Overall, gRNA1, 2, 6, and 10, which exhibited high mKate expression levels (>200% of EF1α promoter), all had 50-60% GC content in gRNA seed sequences, especially within seed 8-10 bp.

**Supplemental Figure 3.**
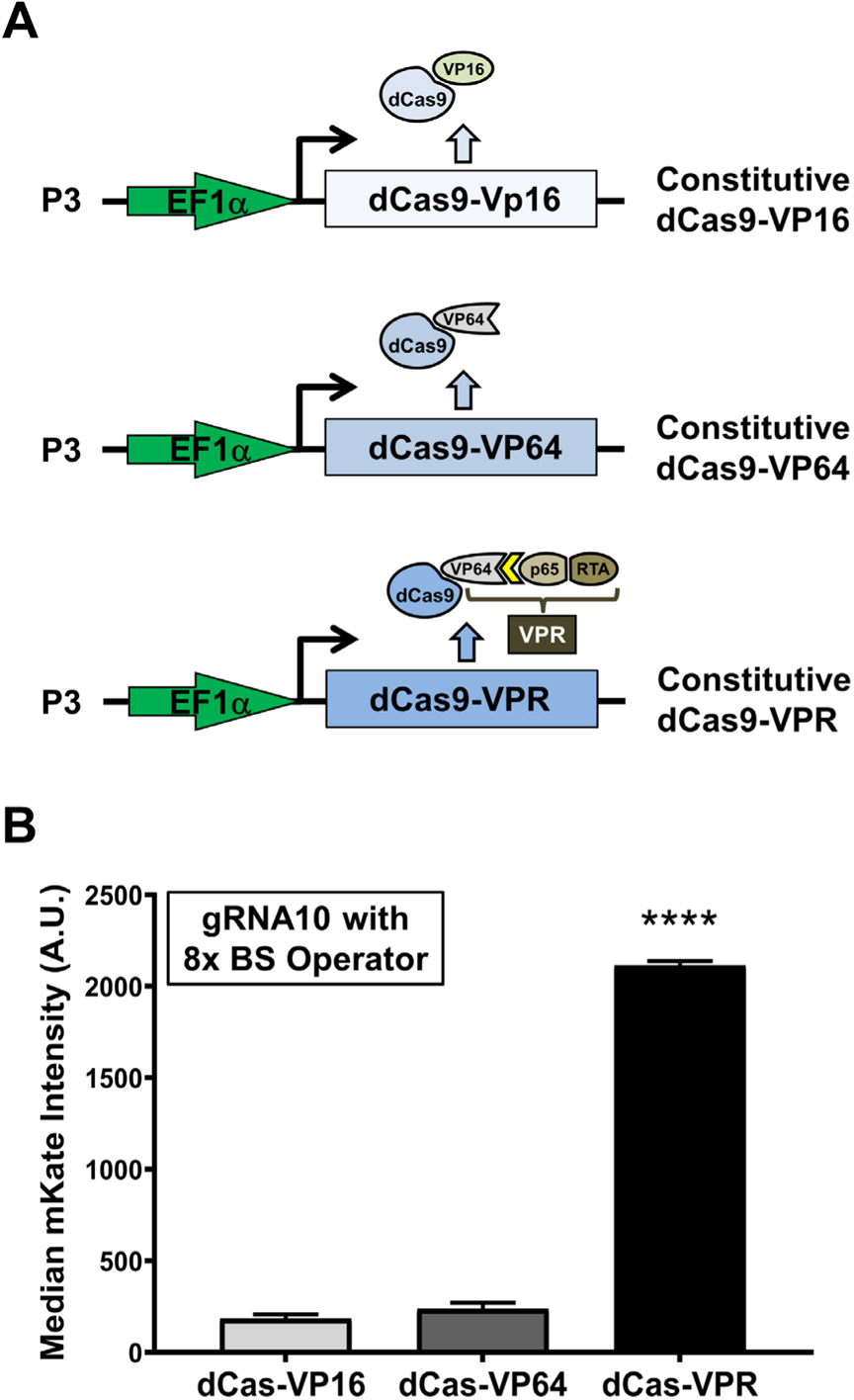
Comparison of gene expression levels obtained with 3 crisprTFs. mKate expression levels were compared for three crisprTFs: dCas-VP16, dCas-VP64, and dCas-VPR, with gRNA10 (P1) and its 8x BS synthetic operator (P2) as depicted in **Figure 1C**. **(A)** Schematic illustration of three versions of plasmid #3 (P3) respectively encoding the three crisprTFs composed of deactivated SpCas9 (dCas9) and transcriptional activation domains (TADs): dCas-VP16, dCas-VP64, and dCas-VPR. **(B)** FACS results showed that dCas-VPR had markedly higher median mKate expression than dCas-VP16 (11.5 fold, *p*<0.0001) and dCas-VP64 (9 fold, *p*<0.0001). Data were presented as median mKate intensity of the entire transfected population with artificial units (A.U.). Data represent the mean ± SD (n = 3) (one-way ANOVA with multiple comparisons corrected by Dunnett test; \*\*\*\**p*<0.0001).

**Supplemental Figure 4.**
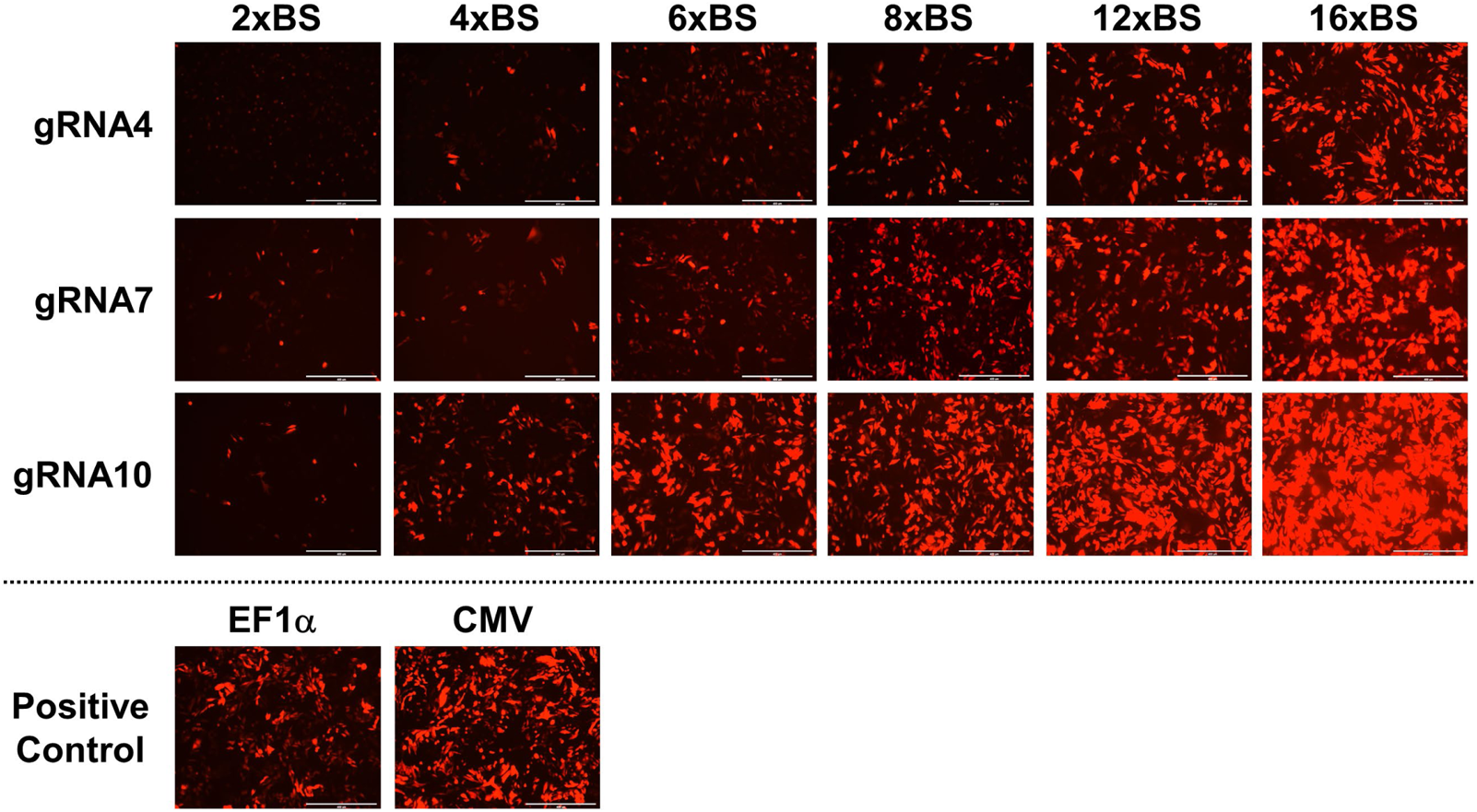
Comparison of gene expression levels with 6 distinct synthetic operators containing different numbers of gRNA BS in three gRNA series. CHO-K1 cells were transfected as illustrated in Figure 1C, with each gRNA constitutively expressed by the U6 promoter from Plasmid #1 (P1) and mKate expressed by each synthetic operator from Plasmid #2 (P2). mKate expression driven by EF1α and CMV promoters served as positive controls. Representative fluorescent images showed a dramatic range of mKate expression among 6 synthetic operators in all three gRNA series at 48 hours post-transfection, especially in the gRNA10 series.

**Supplemental Figure 5.**
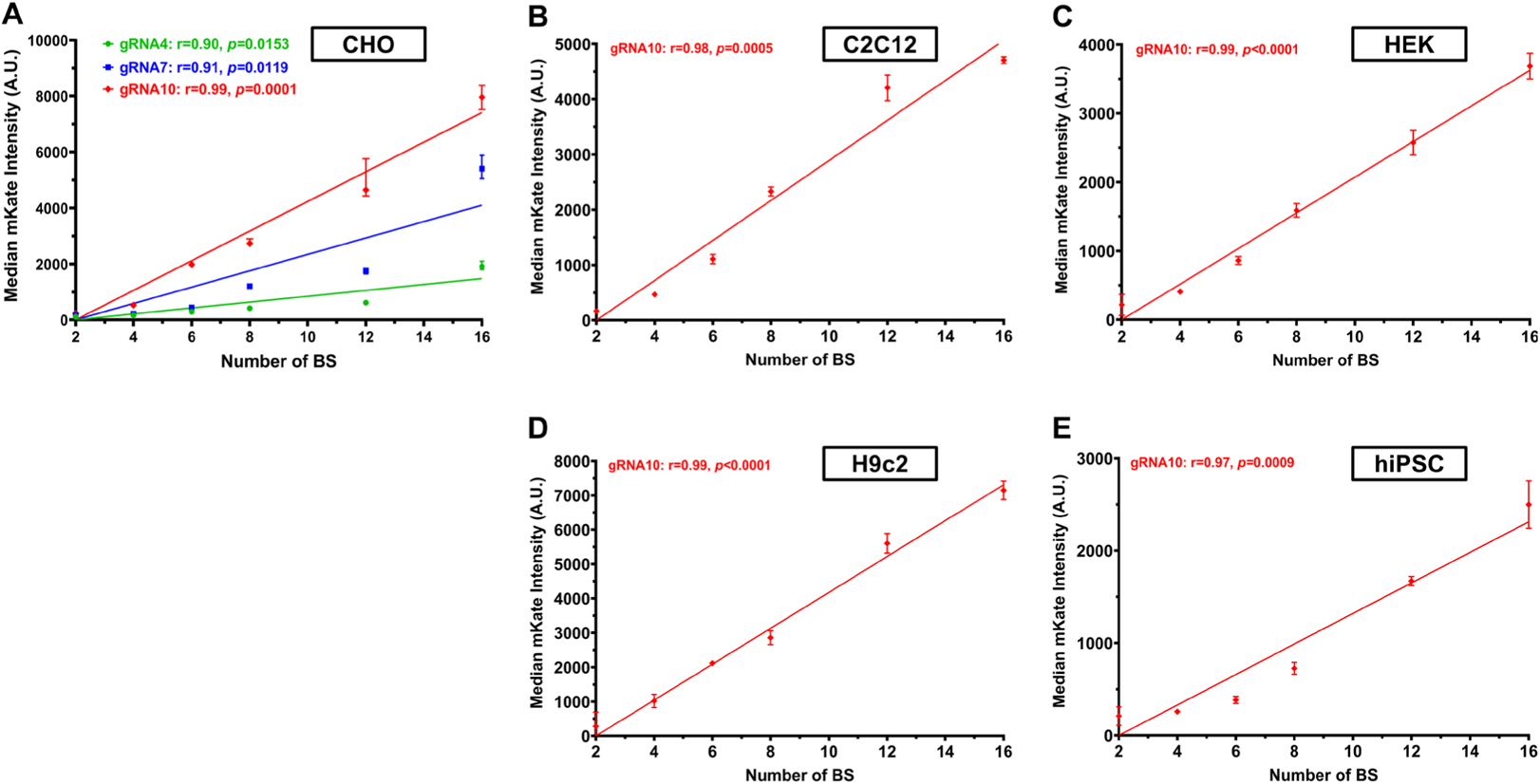
Correlation between the number of gRNA BS in the synthetic operator and the gene expression level. Based on the quantitative results from flow cytometry analyses presented in **Figure 2**, we performed Pearson correlation analysis to reveal the relationship between the number of gRNA BS in each synthetic operator of the three gRNA series and its target gene expression level. **(A)** In CHO-K1 cells, the gRNA4 series had the Pearson correlation coefficient (r)=0.90 (R^2^=0.80, *p*=0.0153); the gRNA7 series had r=0.91 (R^2^=0.83, *p*=0.0119); and the gRNA10 series had r=0.99 (R^2^=0.98, *p*=0.0001). **(B-E)** For the gRNA10 series, r=0.98 (R^2^=0.96, *p*=0.0005) in mouse C2C12 myoblasts (**B**), r=0.99 (R^2^=0.99, *p*<0.0001) in human HEK293T cells (**C**), r=0.99 (R^2^=0.99, *p*<0.0001) in rat H9C2 cardiomyoblast cells (**D**), and r=0.98 (R^2^=0.95, *p*=0.0009) in hiPSC cells (**E**). Simple linear regression was performed to plot the graphs. Data represent the mean ± SD (n = 3).

**Supplemental Figure 6.**
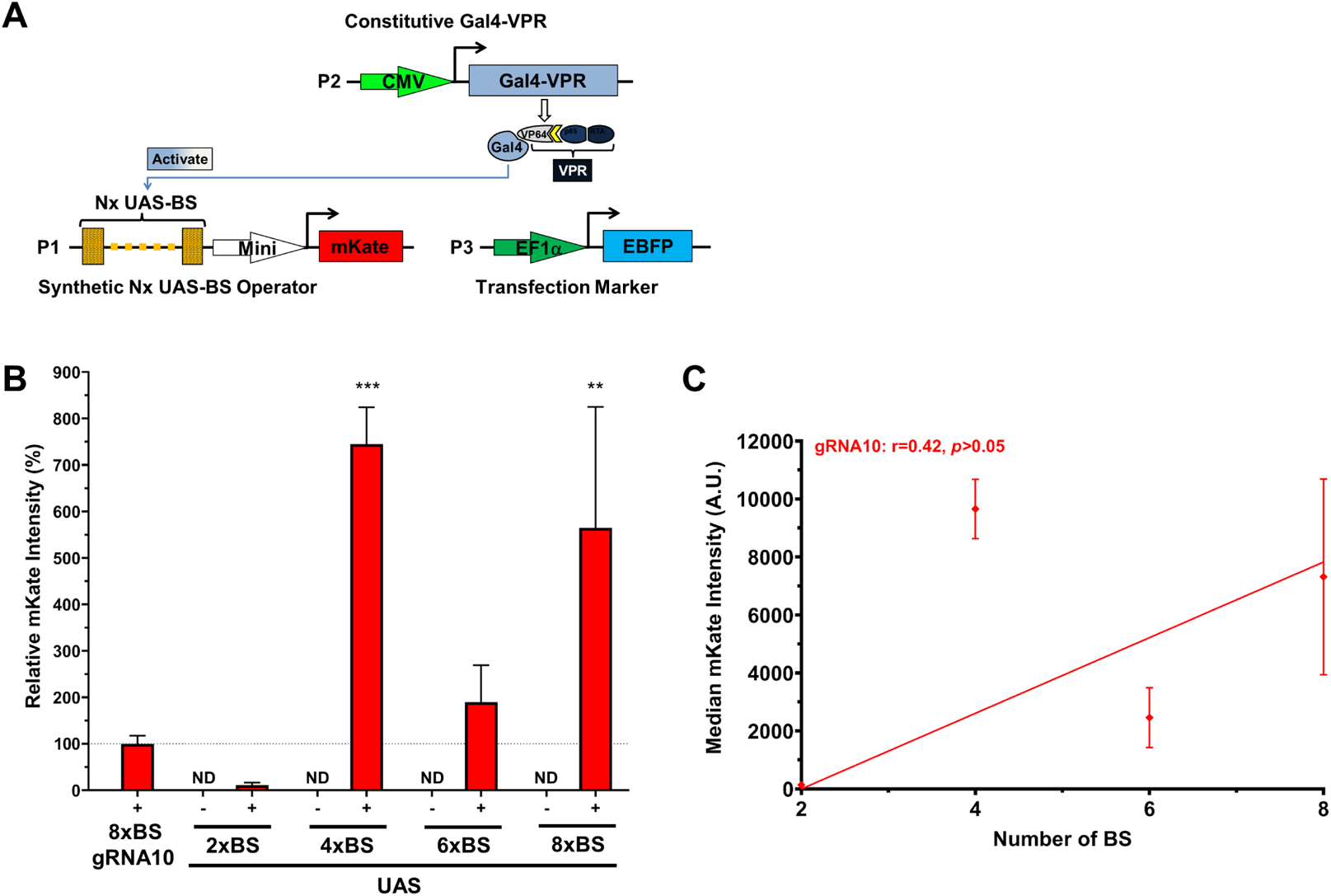
Gene expression controlled by Gal4-VPR/UAS system in CHO-K1 cells. (**A**) A schematic illustration of three plasmids used to transiently transfect CHO-K1 cells: plasmid #1 (P1) encoding the synthetic operator with 2-8x of UAS-BS to drive mKate expression; plasmid #2 (P2) constitutively expressing a Gal4-VPR gene; and plasmid #3 (P3) constitutively expressing the transfection marker (EBFP). (**B**) The mKate signal intensities of representative circuits with 2x-8x UAS-BS. Experimental groups (+), represented by red solid bars, were transfected with three plasmids (P1-P3). Control groups (−), represented by paired red hollow bars, were transfected without P2 (only P1 and P3) to detect baseline UAS operator leakage. (**C**) Correlation between the number of UAS-BS in the synthetic operator and the gene expression level. Data represent the mean ± SD (n = 3) (one-way ANOVA with multiple comparisons corrected by Dunnett test; \*\**p*<0.01, \*\*\**p*<0.001; ND: not detected).

**Supplemental Figure 7.**
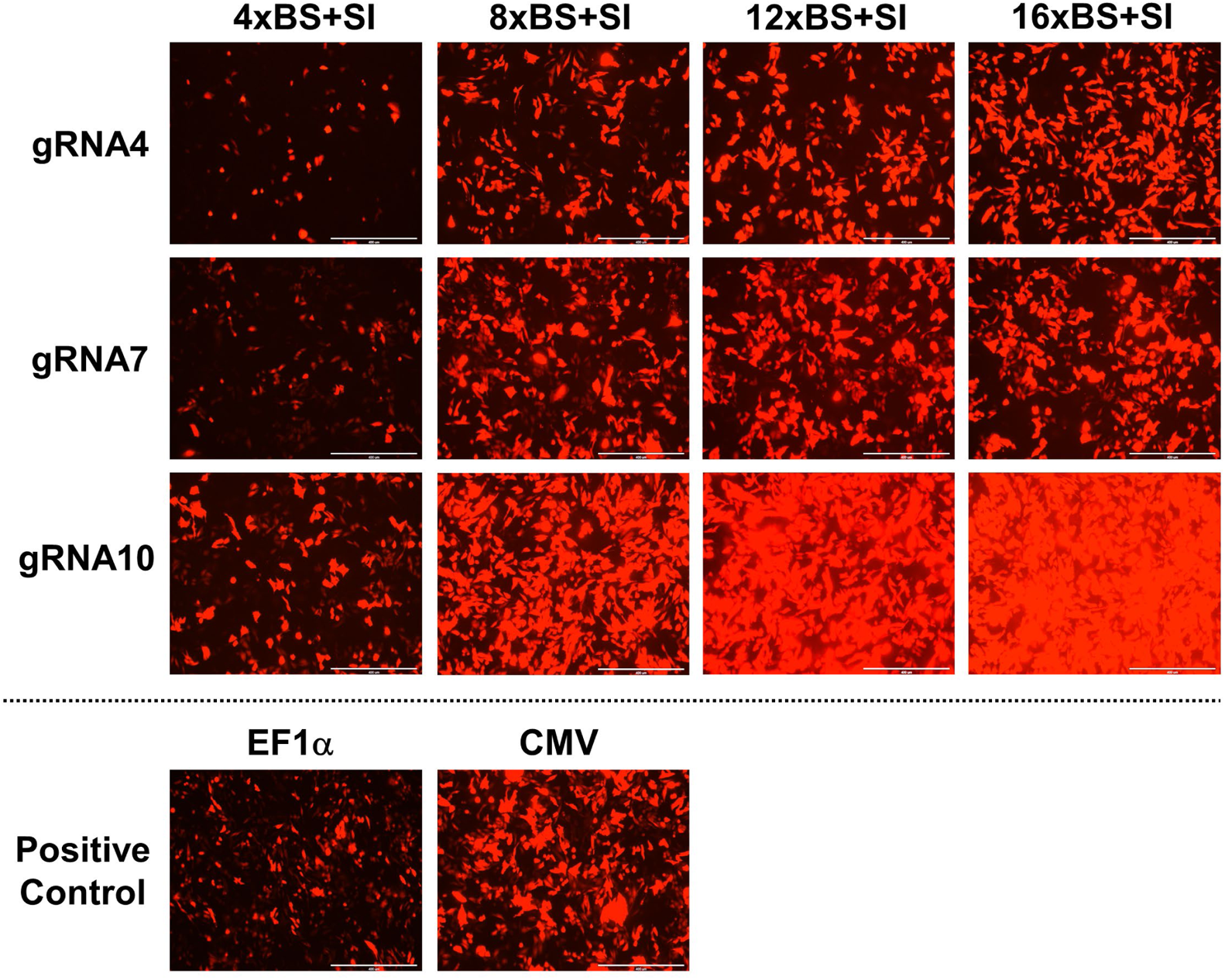
Comparison of gene expression levels with the addition of a synthetic intron (SI) in 4 distinct synthetic operators in three gRNA series. CHO-K1 cells were transfected as illustrated in Figures 1C and 3D, with each gRNA expressed constitutively from P1. mKate was expressed by each synthetic operator (P2) with the presence of an SI at the 5’ UTR of the mKate gene. mKate expression driven by EF1α or CMV promoters served as positive controls. Representative fluorescent images revealed marked increases in mKate expression with the addition of the SI in 4 different synthetic operators of each gRNA series at 48 hours post-transfection. The increment was particularly noteworthy in the gRNA10 series when compared with mKate signals expressed by the same synthetic operators without the SI (data shown in Supplemental Figure 4).

**Supplemental Figure 8.**
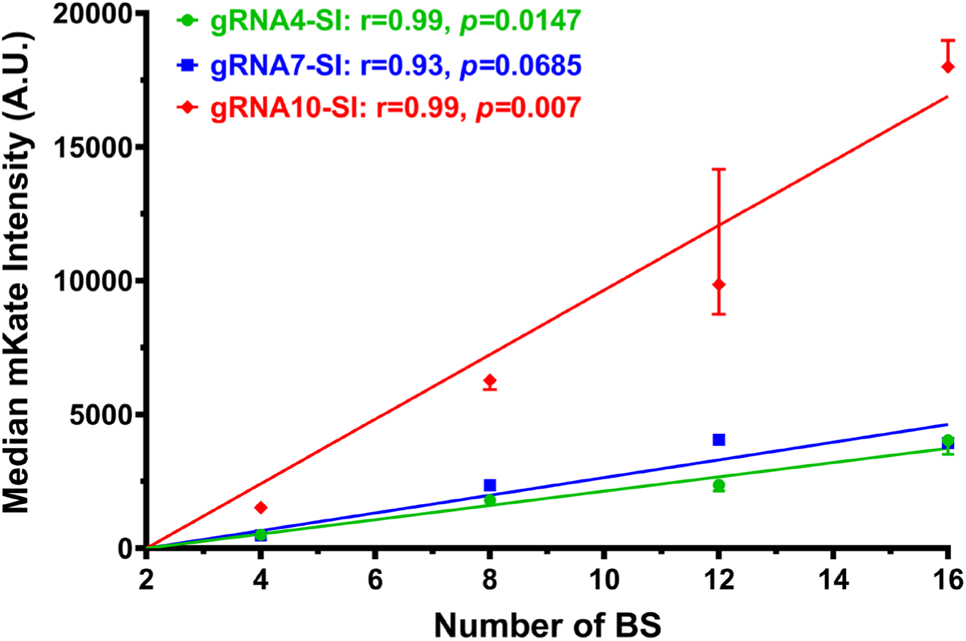
Correlation between the number of gRNA BS and the gene expression level with the addition of a synthetic intron (SI). Pearson correlation analysis uncovered the relationship between the number of gRNA BS in the synthetic operator and the associated gene expression level with the presence of an SI at the 5’ UTR of the target gene. The gRNA4-SI series had the Pearson correlation coefficient (r)=0.99 (R^2^=0.97, *p*=0.0147); the gRNA7-SI series had r=0.93 (R^2^=0.87, *p*=0.0685); and the gRNA10-SI series had r=0.99 (R^2^=0.99, *p*=0.007). Simple linear regression was performed to plot the graph. Data represent the mean ± SD (n = 3).

**Supplemental Figure 9.**
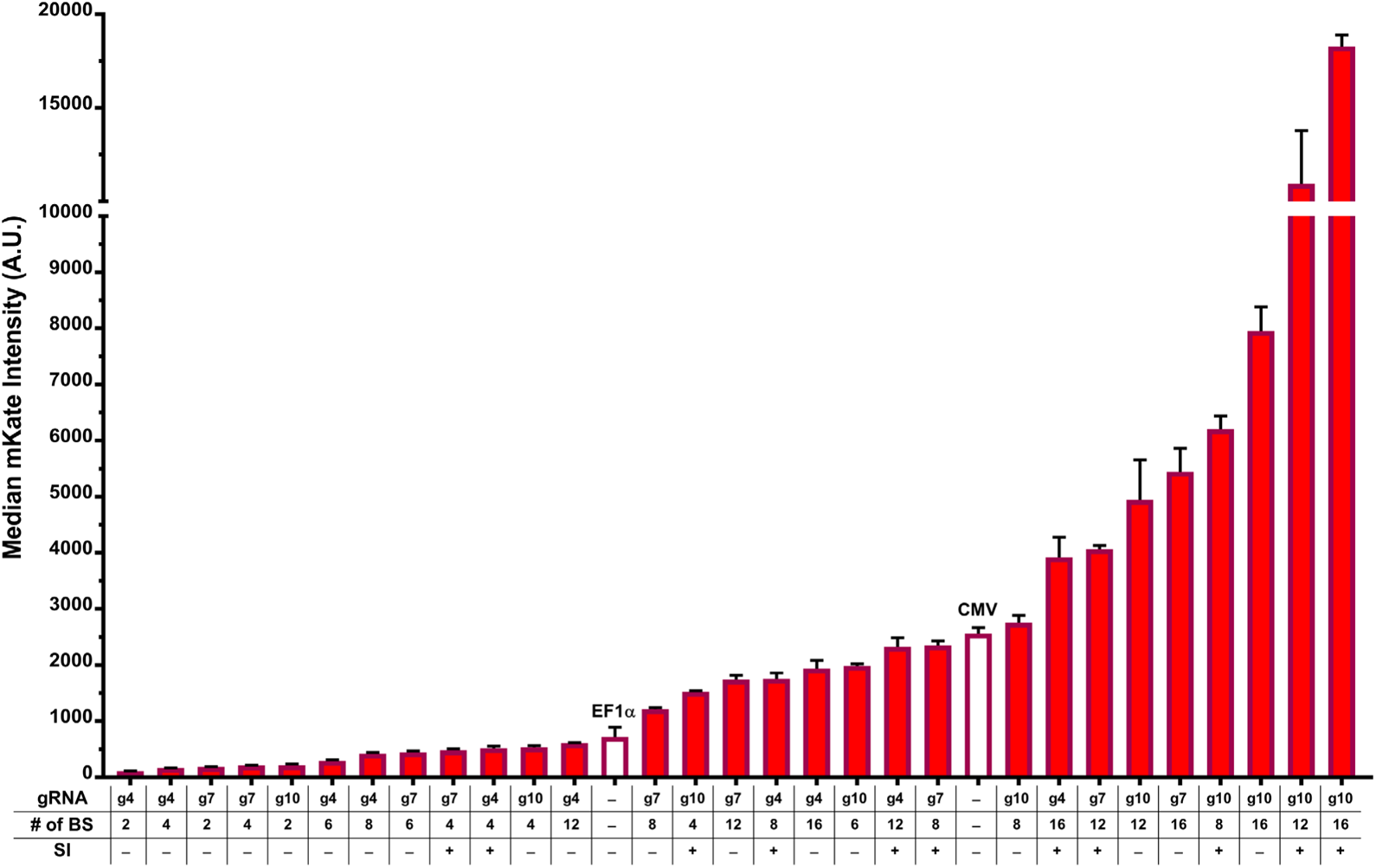
Summary of construct compositions and corresponding gene expression levels. Compositions and gene expression levels of constructs from gRNA4 (g4), gRNA7 (g7), and gRNA10 (g10) series that were tested episomally, with or without the synthetic intron (SI). EF1α and CMV promoter controls are represented by empty bars. Data represent the mean ± SD (n = 3).

**Supplemental Figure 10.**
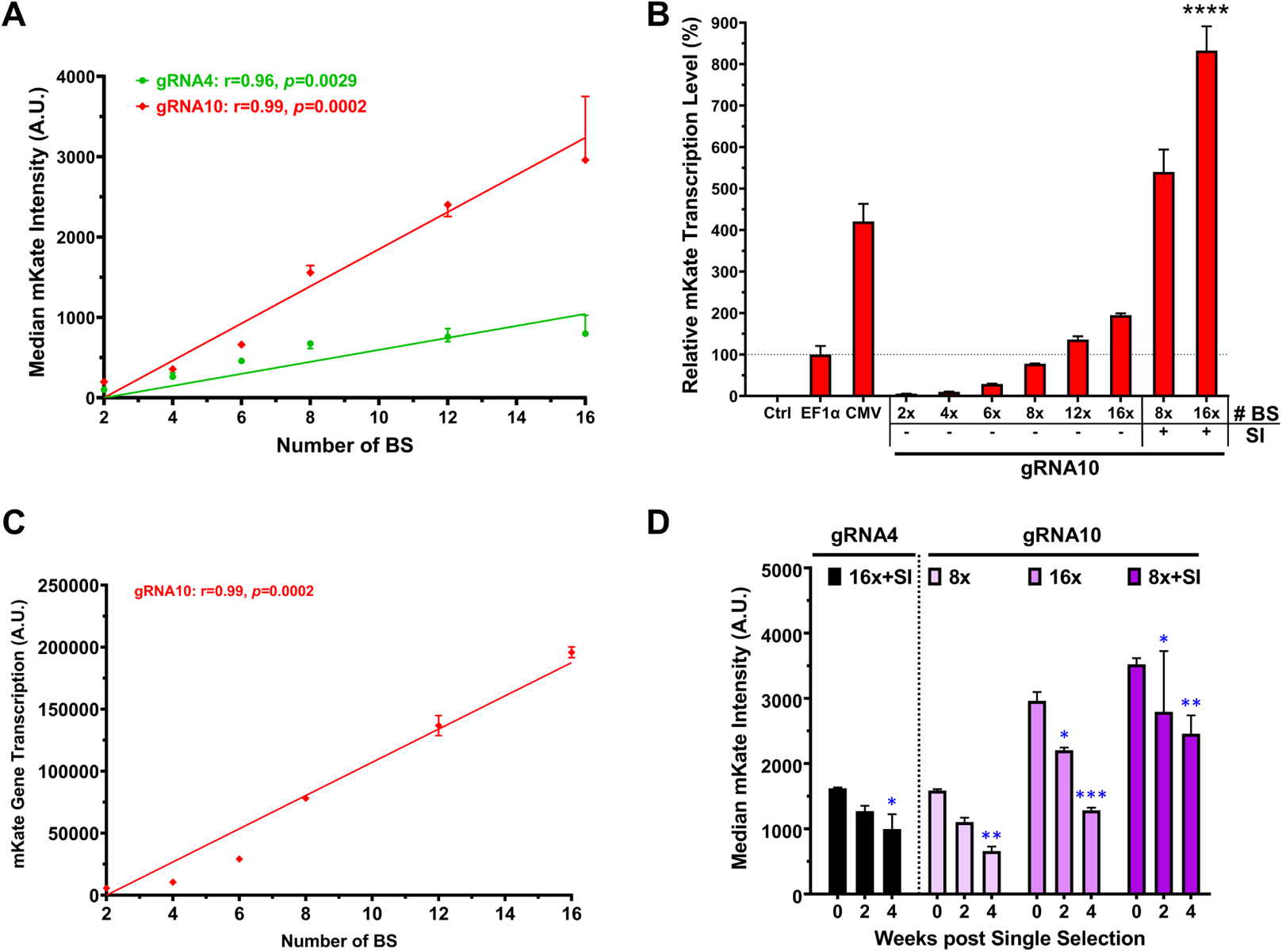
Analyses of chromosomally integrated crisprTF promoter circuits in sLP-CHO cells. **(A)** Pearson correlation analysis showed the relationship between the number of gRNA BS in the synthetic operator and the associated gene expression level when gene circuits were chromosomally integrated. The gRNA4 series had the Pearson correlation coefficient (r)=0.96 (R^2^=0.91, *p*=0.0029), and the gRNA10 series had r=0.99 (R^2^=0.98, *p*=0.0002). Simple linear regression was performed to plot the graph. **(B)** RT-qPCR analysis showing mKate transcription of chromosomally integrated crisprTF promoter circuits relative to the integrated EF1α control (one-way ANOVA with multiple comparisons corrected by Dunnett test). **(C)** Correlation analysis between the number of gRNA10 BS and mKate transcription levels showed r=0.99 (R^2^=0.98, *p*=0.0002). **(D)** The mKate expression levels of representative circuit integrants from gRNA4 (16x BS with SI) and gRNA10 (8x and16x BS without SI, and 8x BS with SI) series over the course of 4 weeks following single antibiotic selection. Data represent the mean ± SD (n = 3) (two-way ANOVA with multiple comparisons corrected by Dunnett test; \**p*≤0.05, \*\**p*≤0.01, \*\*\**p*≤0.001, \*\*\*\**p*≤0.0001).

**Supplemental Figure 11.**
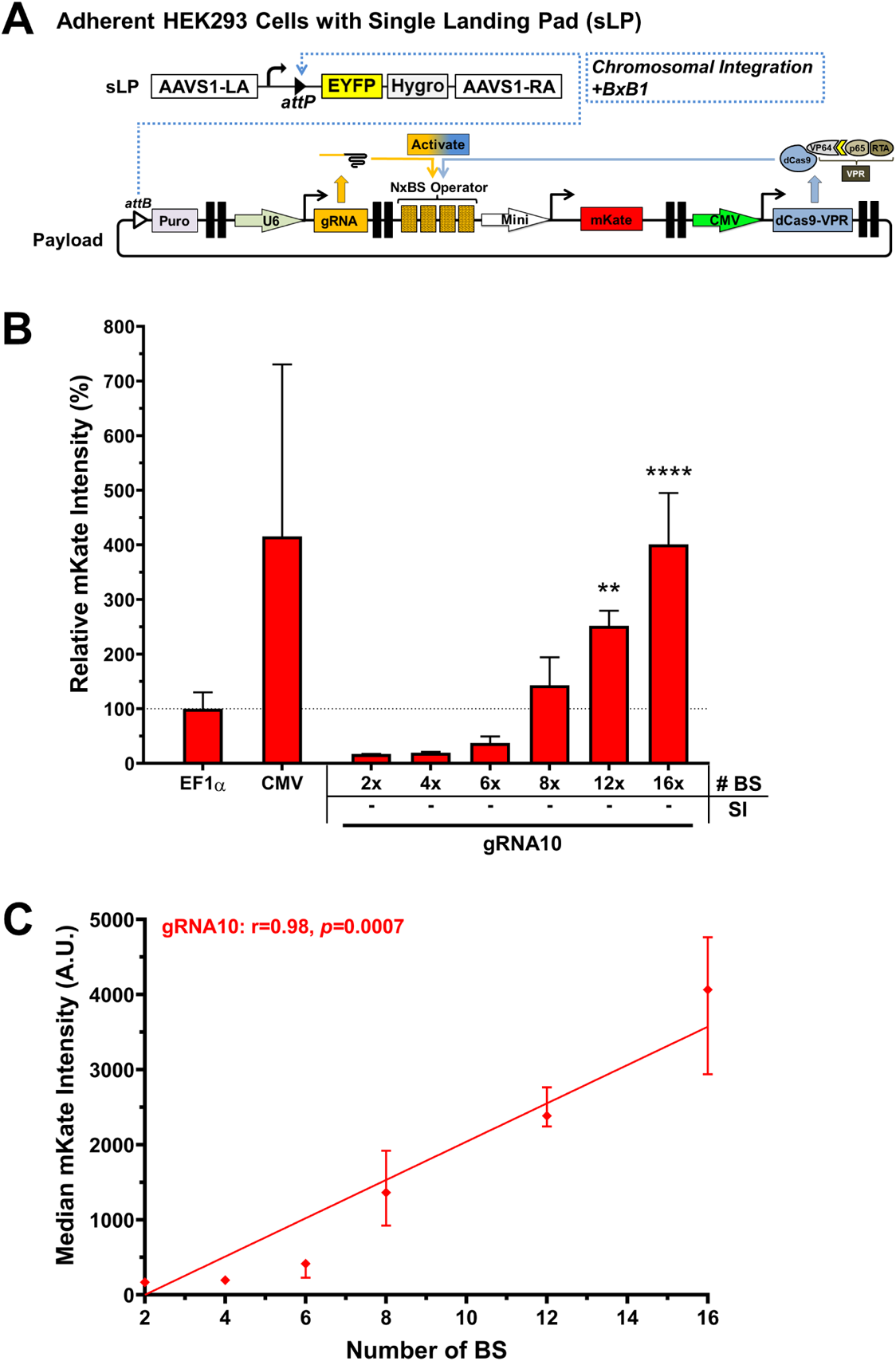
Genomic integration and precision gene expression in HEK293 landing pad cells. **(A)** A schematic illustration of an integration gene circuit and BxB1 recombinase-mediated, site-specific integration in an engineered, adherent HEK293 cell line with a single landing pad (sLP). Positive integration control circuits had a central TU with an EF1α or CMV promoter driving mKate expression and two flanking dummy TUs with no gene expression in the same architecture. **(B)** The mKate signal intensities of the chromosomally integrated payload circuits in sLP-HEK293 cells relative to the integrated EF1α control circuit at 1 week post-selection. **(C)** Correlation between the number of gRNA10 BS and mKate expression levels. Data represent the mean ± SD (n = 3) (one-way ANOVA with multiple comparisons corrected by Dunnett test; \*\**p*<0.01, \*\*\*\**p*<0.0001).

**Supplemental Figure 12.**
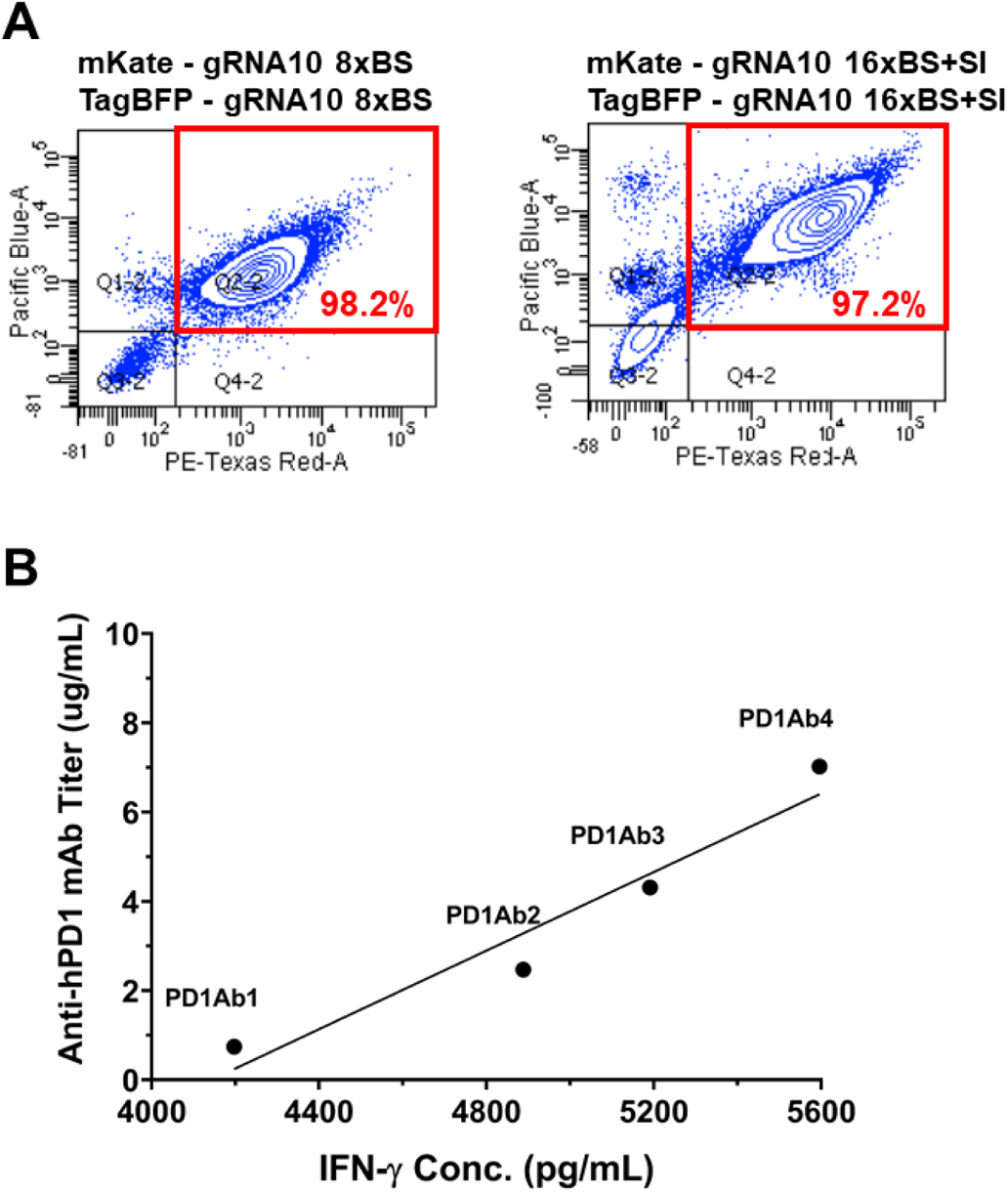
Genomic integration and precision control of target gene expression in the CHO cells engineered with a double landing pad (dLP). **(A)** Precision control of expression levels of two target genes in integrated dLP-CHO cells. Representative FACS dot-plots showing the mKate (x-axis, PE-Texas Red) and TagBFP (y-axis, Pacific Blue) signals in dLP-CHO cells integrated with either 8x BS without SI (left panel) or 16x BS with SI (right panel) control circuit. **(B)** Correlation between IFN-γ production and the anti-hPD1 titer before the start of co-culturing. Pearson correlation analysis revealed the relationship between IFN-γ production and the anti-hPD1 titer in the dLP-CHO cell cultures pre-seeded for 2 days prior to the start of co-culturing (n=3). Pearson correlation coefficient (r)=0.97 (R^2^=0.94, *p*=0.0327).

**Supplemental Figure 13.**
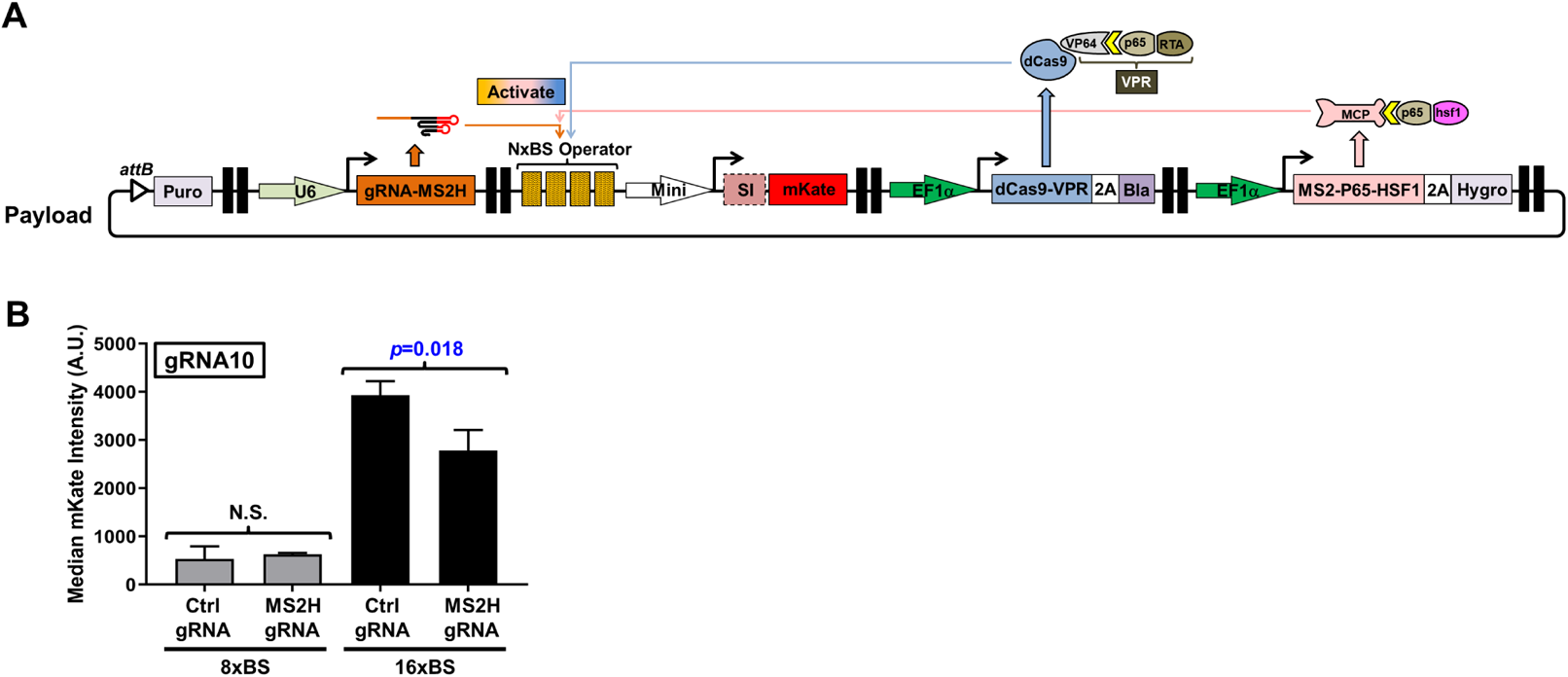
Investigation of the synergistic effect between SAM and dCas-VPR at the chromosomal level. **(A)** A schematic illustration of the gene circuit for chromosomal integration in sLP-CHO cells. The circuit constitutively co-expressed SAM, including gRNA with hairpins and MCP-p65-hsf1, and dCas9-VPR as well as 3 selection marker genes, including the 5’ flanking puromycin and the 3’ flanking blasticidin (associated with dCas9-VPR gene using a self-cleaving P2A peptide) and hygromycin (associated with MCP-p65-hsf1 gene using P2A). **(B)** sLP-CHO cells were transfected with the payload circuit and a BxB1-expressing plasmid. After the triple-antibiotic selection, mKate expression levels assessed by flow cytometry showed no synergistic effect between SAM and dCas-VPR with the chromosomally integrated gRNA10 8xBS operator (*p*>0.05). mKate signals significantly decreased when SAM and dCas-VPR were acting together with the gRNA10 16xBS operator (*p*=0.018). Data represent the mean ± SD (n = 3) (two-tailed paired Student’s *t*-test).

**Supplemental Figure 14.**
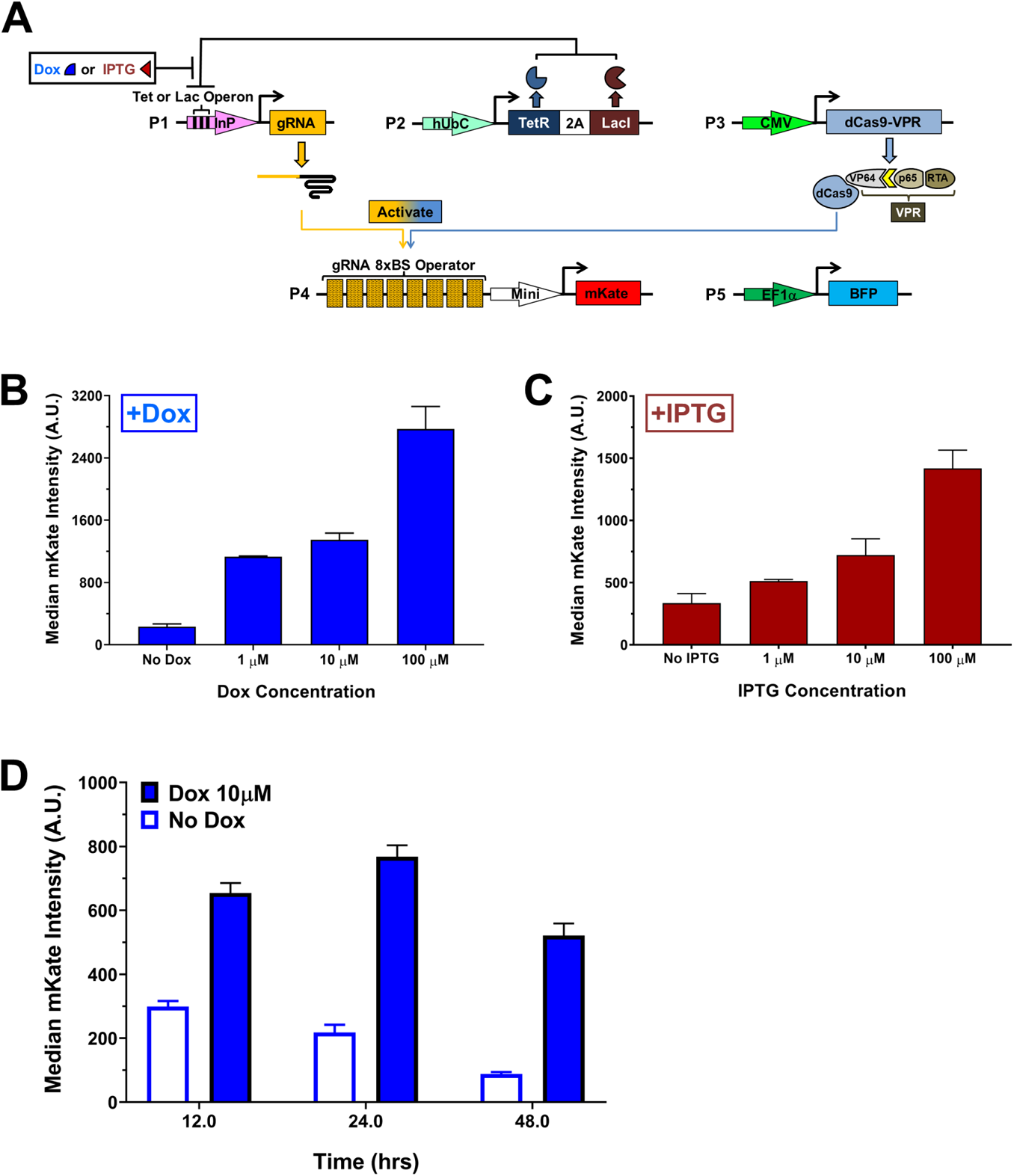
crisprTF promoters with small molecule-inducible gRNA expression. **(A)** To equip the crisprTF promoter system with an added tier of controllability, we developed inducible switches with doxycycline (Dox)-inducible or isopropyl β-_D_-1-thiogalactopyranoside (IPTG)-inducible gRNA expression. Either a Tet or a Lac operon was custom imbedded into a RNA polymerase III (Pol III) promoter driving gRNA10 expression to render inducibility. Without an appropriate small molecule inducer (Dox or IPTG), the Tet repressor or Lac inhibitor (both constitutively expressed from a single plasmid using a self-cleaving P2A peptide) bound to the Tet or Lac operon, respectively, and repressed gRNA expression. In the presence of Dox or IPTG, the Tet repressor or Lac inhibitor did not bind to the respective operon, permitting gRNA10 transcription. **(B)** Titration analysis of the Dox-inducible gRNA10 expression with its 8x BS synthetic promoter unveiled incrementally increased mKate expression with increased Dox concentration; the highest expression level was seen with 100 μM Dox. **(C)** Titration analysis of the IPTG-inducible gRNA10 expression with its 8x BS synthetic promoter revealed incrementally increased mKate expression with increased IPTG concentration; the highest expression level was seen with 100 μM IPTG. **(D)** mKate expression kinetics in the presence (solid blue bars) or absence (empty blue bars) of 10 μM Dox. Data represent the mean ± SD (n = 3).

