## Supplemental Table 1 for "A Synthetic Transcription Platform for Programmable Gene Expression in Mammalian Cells"

**Supplemental Table 1. Sequences of key genetic parts and primers**

| Genetic Part Name | DNA sequence | Appearance in Figure(s) |
|---|---|---|
| 2A sequence | ggatctggcgccaccaacttctctctgctgaagcaggccggcgacgtggaggagaacccaggcccc | 4, 5, 6, S13, S14 |
| 2x cHS4 insulator | aagctttttccccgtatccccccaggtgtctgcaggctcaaagagcagcgagaagcgttcagaggaaagcgatcccgtgccaccttccccgtgcccgggctgtccccgcacgctgccggctcggggatgcggggggagcgccggaccggagcggagccccgggcggctcgctgctgccccctagcgggggagggacgtaattacatccctgggggctttgggagggggctgtccccgtgagctcccaggcgcgcctgtcattctaaatctctctttcagcctaaagctttttccccgtatccccccaggtgtctgcaggctcaaagagcagcgagaagcgttcagaggaaagcgatcccgtgccaccttccccgtgcccgggctgtccccgcacgctgccggctcggggatgcgggggagcgccggaccggagcggagccccgggcggctcgctgctgccccctagcgggggagggacgtaattacatccctgggggctttgggggggggctgtccccgtgagctc | 1, 4, 5, 6, S11, S13 |
| gRNA scaffold | NNNNNNNNNNNNNNNNNNNNgttttagagctagaaatagcaagttaaaataaggctagtccgttatcaacttgaaaaagtggcaccgagtcggtgc | 1-6, S1-S14 |
| sgRNA10 16x BS + mini CMV promoter | tacgagggcgatccaacgactggtacgagggcgatccaacgacgggtacgagggcgatccaacgacaggtacgagggcgatccaacgaccggtacgagggcgatccaacgactggtacgagggcgatccaacgacgggtacgagggcgatccaacgacaggtacgagggcgatccaacgaccggtacgagggcgatccaacgactggtacgagggcgatccaacgacgggtacgagggcgatccaacgacaggtacgagggcgatccaacgaccggtacgagggcgatccaacgactggtacgagggcgatccaacgacgggtacgagggcgatccaacgacaggtacgagggcgatccaacgaccggtggtctatataagcagagctcgtttagtgaaccgtcagatcgcctggagacgccatccacgctgttttgacctccatagaagac | 2-6, S4-S14 |
| sgRNA10 16x BS + mini CMV promoter + synthetic intron | tacgagggcgatccaacgactggtacgagggcgatccaacgacgggtacgagggcgatccaacgacaggtacgagggcgatccaacgaccggtacgagggcgatccaacgactggtacgagggcgatccaacgacgggtacgagggcgatccaacgacaggtacgagggcgatccaacgaccggtacgagggcgatccaacgactggtacgagggcgatccaacgacgggtacgagggcgatccaacgacaggtacgagggcgatccaacgaccggtacgagggcgatccaacgactggtacgagggcgatccaacgacgggtacgagggcgatccaacgacaggtacgagggcgatccaacgaccggtggtctatataagcagagctcgtttagtgaaccgtcagatcgcctggagacgccatccacgctgttttgacctccatagaagacgtgagtactccctctcaaaagcgggcatgacttctgcgctaagattgtcagtttccaaaaacgaggaggatttgatattcacctggcccgcggtgatgcctttgagggtggccgcgtccatctggtcagaaaagacaatctttttgttgtcaagcttgaggtgtggcaggcttgagatctggccatacacttgagtgacaatgacatccactttgcctttctctccacag | 3-6, S7-S10, S12-S13 |
| sgRNA4 16x BS + mini CMV promoter | cgcgagggtagttgacaacgtggcgcgagggtagttgacaacgggcgcgagggtagttgacaacgaggcgcgagggtagttgacaacgcggcgcgagggtagttgacaacgtggcgcgagggtagttgacaacgggcgcgagggtagttgacaacgaggcgcgagggtagttgacaacgcggcgcgagggtagttgacaacgtggcgcgagggtagttgacaacgggcgcgagggtagttgacaacgaggcgcgagggtagttgacaacgcggcgcgagggtagttgacaacgtggcgcgagggtagttgacaacgggcgcgagggtagttgacaacgaggcgcgagggtagttgacaacgcggtggtctatataagcagagctcgtttagtgaaccgtcagatcgcctggagacgccatccacgctgttttgacctccatagaagac | 2, 3, 4, S4, S5, S7-S10 |
| sgRNA7 16x BS + mini CMV promoter | agtacgggtcgtacacgaagtggagtacgggtcgtacacgaaggggagtacgggtcgtacacgaagaggagtacgggtcgtacacgaagcggagtacgggtcgtacacgaagtggagtacgggtcgtacacgaaggggagtacgggtcgtacacgaagaggagtacgggtcgtacacgaagcggagtacgggtcgtacacgaagtggagtacgggtcgtacacgaaggggagtacgggtcgtacacgaagaggagtacgggtcgtacacgaagcggagtacgggtcgtacacgaagtggagtacgggtcgtacacgaaggggagtacgggtcgtacacgaagaggagtacgggtcgtacacgaagcggtggtctatataagcagagctcgtttagtgaaccgtcagatcgcctggagacgccatccacgctgttttgacctccatagaagac | 2, 3, S4, S5, S7-S9 |
| U6 promoter + gRNA10 + MS2 Stem Loop RNA | agagggcctatttcccatgattccttcatatttgcatatacgatacaaggctgttagagagataattagaattaatttgactgtaaacacaaagatattagtacaaaatacgtgacgtagaaagtaataatttcttgggtagtttgcagttttaaaattatgttttaaaatggactatcatatgcttaccgtaacttgaaagtatttcgatttcttggctttatatatcttgtggaaaggacgaaacaccgtacgagggcgatccaacgagttttagagctaggccaacatgaggatcacccatgtctgcagggcctagcaagttaaaataaggctagtccgttatcaacttggccaacatgaggatcacccatgtctgcagggccaagtggcaccgagtcggtgctttttttctag | 3, S13 |
| VPR (VP64-SV40NLS-p65-RTA) | gaggccagcggttccggacgggctgacgcattggacgattttgatctggatatgctgggaagtgacgccctcgatgattttgaccttgacatgcttggttcggatgcccttgatgactttgacctcgacatgctcggcagtgacgcccttgatgatttcgacctggacatgctgattaactctagaagttccggatctccgaaaaagaaacgcaaagttggtagccagtacctgcccgacaccgacgaccggcaccggatcgaggaaaagcggaagcggacctacgagacattcaagagcatcatgaagaagtccccttcagcggccccaccgaccctagacctccacctagaagaatcgccgtgcccagcagatccagcgccagcgtgccaaaacctgccccccagccttaccccttcaccagcagcctgagcaccatcaactacgacgagttccctaccatggtgttccccagcggccagatctctcaggcctctgctctggctccagcccctcctcaggtgctgcctcaggctcctgctcctgcaccagctccagccatggtgtctgcactggctcaggcaccagcacccgtgcctgtgctggctcctggacctccacaggctgtggctccaccagcccctaaacctacacaggccggcgagggcacactgtctgaagctctgctgcagctgcagttcgacgacgaggatctgggagccctgctgggaaacagcaccgatcctgccgtgttcaccgacctggccagcgtggacaacagcgagttccagcagctgctgaaccagggcatccctgtggcccctcacaccaccgagcccatgctgatggaataccccgaggccatcacccggctcgtgacaggcgctcagaggcctcctgatccagctcctgcccctctgggagcaccaggcctgcctaatggactgctgtctggcgacgaggacttcagctctatcgccgatatggatttctcagccttgctgggctctggcagcggcagccgggattccagggaagggatgtttttgccgaagcctgaggccggctccgctattagtgacgtgtttgagggccgcgaggtgtgccagccaaaacgaatccggccatttcatcctccaggaagtccatgggccaaccgcccactccccgccagcctcgcaccaacaccaaccggtccagtacatgagccagtcgggtcactgaccccggcaccagtccctcagccactggatccagcgcccgcagtgactcccgaggccagtcacctgttggaggatcccgatgaagagacgagccaggctgtcaaagcccttcgggagatggccgatactgtgattccccagaaggaagaggctgcaatctgtggccaaatggacctttcccatccgcccccaaggggccatctggatgagctgacaaccacacttgagtccatgaccgaggatctgaacctggactcacccctgaccccggaattgaacgagattctggataccttcctgaacgacgagtgcctcttgcatgccatgcatatcagcacaggactgtccatcttcgacacatctctgttttga | 1-6, S1, S3-S5, S7-S12 |
| G.Luc sp CDS Signal peptide + Anti-huPD1 (5C4) Light Chain | ggagttaaggtattgttcgcattgatttgcatagccgtggccgaagctgaaatagtgctgacccagtcccccgccaccctctctctctcacccggagagagggcaactctgtcctgcagggcctcccaatccgtaagcagctatctggcttggtaccagcaaaaacccggacaggcaccccggctgttgatctatgacgcctccaaccgagccactggcatccccgccagattctccggatctggatcagggactgattttaccctgactatcagttctctggaacctgaagattttgcagtttattattgtcagcaaagctctaactggccaagaacctttggccagggcacgaaggtcgagataaaaactgtcgcggccccaagcgtctttatctttccgccgtctgatgaacagctgaaatcagggaccgcctctgtggtgtgtctgctgaacaacttctacccaagggaagctaaggtgcaatggaaagtagacaatgccttgcagagcggcaactctcaggagagcgtaaccgaacaggatagtaaggattcaacttacagtctgagctctaccctgactctgagtaaggccgattacgagaagcataaagtctacgcatgcgaggtcacacatcaaggtctttcatctcccgtgacaaagtcctttaatcgaggggaatgctaa | 6 |

| | | |
|---|---|---|
| G.Luc sp Signal peptide + Anti-huPD1 (5C4) Heavy Chain | atgggagttaaggtattgttcgcattgatttgcatagccgtggccgaagctcaggtgcagctcgtagagagcgggggaggtgtcgttcagccagggaggagtctgcggctggactgcaaagcctccgggattactttctcaaacagcgggatgcactgggtgaggcaggctcccggaaagggcctcgagtgggtggcggtaatatggtacgacgggagcaaacgctactacgcagacagtgtgaaaggaaggttcactatttctcgcgacaattcaaagaacaccctcttcctgcagatgaacagtctgcgggcagaagacaccgctgtctattactgtgccaccaatgacgattactggggccagggtaccctcgtgaccgtgtctagcgcctctaccaaaggaccatcagtttttcctcttgctccctgcagtcgctcaaccagcgagagcacagcagcgctgggatgtctggtgaaggattattttcctgagcccgtgactgtgagctggaattcaggcgccctgacgtccggcgtccatacattcccagcggtacttcaaagtagcgggttgtactctctctctagcgtggtaaccgtaccgagctcctccctggggacgaaaacgtatacatgtaatgtcgatcacaaaccatctaacacaaaagtggacaaacgcgttgagtccaagtatggccctccatgcccaccctgccccgcaccggagtttctgggcgggcccagtgtctttctgttcccaccgaagcctaaggacacgttgatgatctcaagaacacctgaagtcacctgcgtagtcgtggacgtttctcaggaggatcccgaggtccaattcaattggtacgttgatggagtggaggtccacaacgcaaagacaaagccgcgcgaagaacagttcaattctacttaccgcgttgttagcgtgctgactgtgctccaccaagactggctgaatggtaaggagtataagtgcaaggtgagcaataagggattgccatctagcatcgaaaagacaatatccaaagccaagggccaaccacgagagccacaagtgtacacgttgcctccctcacaagaagagatgaccaagaatcaagtgagcctcacttgcctggtcaagggattctacccttctgatatcgcagtggagtgggagtccaatggacagcccgagaacaactacaagacaacacccccagtgctggattccgacggctcattcttcttgtatagccggctgacagtggacaagagcaggtggcaggaaggaaatgtcttctcctgctccgtgatgcacgaggccctccacaaccactacactcagaaatctctctctctttcactcggtaaataa | 6 |
| Gal4 DNA binding domain | atgaagctactgtcttctatcgaacaagcatgcgatatttgccgacttaaaaagctcaagtgctccaaagaaaaaccgaagtgcgccaagtgtctgaagaacaactgggagtgtcgctactctcccaaaaccaaaaggtctccgctgactagggcacatctgacagaagtggaatcaaggctagaaagactggaacagctatttctactgatttttcctcgagaagaccttgacatgattttgaaaatggattctttacaggatataaaagcattgttaacaggattatttgtacaagataatgtgaataaagatgccgtcacagatagattggcttcagtggagactgatatgcctctaacattgagacagcatagaataagtgcgacatcatcatcggaagagagtagtaacaaaggtcaaagacagttgactgtatcg | S6 |
| 2x UAS for Gal4DBD binding + mini CMV promoter | cggagtactgtcctccgagcggagtactgtcctccgagtggtctatataagcagagctcgtttagtgaaccgtcagatcgcctggagacgccatccacgctgttttgacctccatagaagac | S6 |
| 4x UAS for Gal4DBD binding + mini CMV promoter | cggagtactgtcctccgagcggagtactgtcctccgagcggagtactgtcctccgagcggagtactgtcctccgagtggtctatataagcagagctcgtttagtgaaccgtcagatcgcctggagacgccatccacgctgttttgacctccatagaagac | S6 |
| 6x UAS for Gal4DBD binding + mini CMV promoter | cggagtactgtcctccgagcggagtactgtcctccgagcggagtactgtcctccgagcggagtactgtcctccgagcggagtactgtcctccgagcggagtactgtcctccgagtggtctatataagcagagctcgtttagtgaaccgtcagatcgcctggagacgccatccacgctgttttgacctccatagaagac | S6 |
| 8x UAS for Gal4DBD binding + mini CMV promoter | cggagtactgtcctccgagcggagtactgtcctccgagcggagtactgtcctccgagcggagtactgtcctccgagcggagtactgtcctccgagcggagtactgtcctccgagcggagtactgtcctccgagcggagtactgtcctccgagtggtctatataagcagagctcgtttagtgaaccgtcagatcgcctggagacgccatccacgctgttttgacctccatagaagac | S6 |

| Primers | DNA sequence |
|---|---|
| RT-mKate-F | GGTGAACTTCCCATCCAACG |
| RT-mKate-R | ATGTCGCTTCTGCCTTCCAG |
| RT-GAPDH-F | GCACCACCAACTGCTTAGCC |
| RT-GAPDH-R | GGGCCATCCACAGTCTTCTG |
