## Supplemental Table 2 for "A Synthetic Transcription Platform for Programmable Gene Expression in Mammalian Cells"

**Supplemental Table 2. List of plasmids used in experiments.**

| Name | Description | Purpose | First Appearance in Figure |
|---|---|---|---|
| pWC0011 | pU6-sgRNA1 | constitutively expressing gRNA1 | 1 |
| pWC0012 | pU6-sgRNA2 | constitutively expressing gRNA2 | 1 |
| pWC0013 | pU6-sgRNA3 | constitutively expressing gRNA3 | 1 |
| pWC0014 | pU6-sgRNA4 | constitutively expressing gRNA4 | 1 |
| pWC0015 | pU6-sgRNA5 | constitutively expressing gRNA5 | 1 |
| pWC0016 | pU6-sgRNA6 | constitutively expressing gRNA6 | 1 |
| pWC0017 | pU6-sgRNA7 | constitutively expressing gRNA7 | 1 |
| pWC0018 | pU6-sgRNA8 | constitutively expressing gRNA8 | 1 |
| pWC0019 | pU6-sgRNA9 | constitutively expressing gRNA9 | 1 |
| pWC0020 | pU6-sgRNA10 | constitutively expressing gRNA10 | 1 |
| pWC0021 | p8xBS (sgRNA1)-mini-mKate | encoding the synthetic operator with 8 of gRNA1 BS to drive mKate expression | 1 |
| pWC0022 | p8xBS (sgRNA2)-mini-mKate | encoding the synthetic operator with 8 of gRNA2 BS to drive mKate expression | 1 |
| pWC0023 | p8xBS (sgRNA3)-mini-mKate | encoding the synthetic operator with 8 of gRNA3 BS to drive mKate expression | 1 |
| pWC0024 | p8xBS (sgRNA5)-mini-mKate | encoding the synthetic operator with 8 of gRNA5 BS to drive mKate expression | 1 |
| pWC0025 | p8xBS (sgRNA6)-mini-mKate | encoding the synthetic operator with 8 of gRNA6 BS to drive mKate expression | 1 |
| pWC0026 | p8xBS (sgRNA8)-mini-mKate | encoding the synthetic operator with 8 of gRNA8 BS to drive mKate expression | 1 |
| pWC0027 | p8xBS (sgRNA9)-mini-mKate | encoding the synthetic operator with 8 of gRNA9 BS to drive mKate expression | 1 |
| pWC0030 | pEF1a-dCas9-VP16 | constitutively expressing a dCas9-VP16 | S3 |
| pWC0031 | pEF1a-dCas9-VP64 | constitutively expressing a dCas9-VP64 | S3 |
| pWC0032 | pEF1a-dCas9-VPR | constitutively expressing a dCas9-VPR | S3 |
| pWC0033 | pCMV-dCas9-VPR | constitutively expressing a crisprTF | 1 |
| pWC0034 | p2xBS (sgRNA4)-mini-mKate | encoding the synthetic operator with 2 of gRNA4 BS to drive mKate expression | 2 |
| pWC0035 | p4xBS (sgRNA4)-mini-mKate | encoding the synthetic operator with 4 of gRNA4 BS to drive mKate expression | 2 |
| pWC0036 | p6xBS (sgRNA4)-mini-mKate | encoding the synthetic operator with 6 of gRNA4 BS to drive mKate expression | 2 |
| pWC0037 | p8xBS (sgRNA4)-mini-mKate | encoding the synthetic operator with 8 of gRNA4 BS to drive mKate expression | 1 |
| pWC0038 | p12xBS (sgRNA4)-mini-mKate | encoding the synthetic operator with 12 of gRNA4 BS to drive mKate expression | 2 |
| pWC0039 | p16xBS (sgRNA4)-mini-mKate | encoding the synthetic operator with 16 of gRNA4 BS to drive mKate expression | 2 |
| pWC0040 | p2xBS (sgRNA7)-mini-mKate | encoding the synthetic operator with 2 of gRNA7 BS to drive mKate expression | 2 |
| pWC0041 | p4xBS (sgRNA7)-mini-mKate | encoding the synthetic operator with 4 of gRNA7 BS to drive mKate expression | 2 |
| pWC0042 | p6xBS (sgRNA7)-mini-mKate | encoding the synthetic operator with 6 of gRNA7 BS to drive mKate expression | 2 |
| pWC0043 | p8xBS (sgRNA7)-mini-mKate | encoding the synthetic operator with 8 of gRNA7 BS to drive mKate expression | 1 |
| pWC0044 | p12xBS (sgRNA7)-mini-mKate | encoding the synthetic operator with 12 of gRNA7 BS to drive mKate expression | 2 |
| pWC0045 | p16xBS (sgRNA7)-mini-mKate | encoding the synthetic operator with 16 of gRNA7 BS to drive mKate expression | 2 |
| pWC0046 | p2xBS (sgRNA10)-mini-mKate | encoding the synthetic operator with 2 of gRNA10 BS to drive mKate expression | 2 |
| pWC0047 | p4xBS (sgRNA10)-mini-mKate | encoding the synthetic operator with 4 of gRNA10 BS to drive mKate expression | 2 |
| pWC0048 | p6xBS (sgRNA10)-mini-mKate | encoding the synthetic operator with 6 of gRNA10 BS to drive mKate expression | 2 |
| pWC0049 | p8xBS (sgRNA10)-mini-mKate | encoding the synthetic operator with 8 of gRNA10 BS to drive mKate expression | 1 |
| pWC0050 | p12xBS (sgRNA10)-mini-mKate | encoding the synthetic operator with 12 of gRNA10 BS to drive mKate expression | 2 |
| pWC0051 | p16xBS (sgRNA10)-mini-mKate | encoding the synthetic operator with 16 of gRNA10 BS to drive mKate expression | 2 |
| pWC0052 | pEF1a-EBFP | constitutively expressing the transfection marker (EBFP) | 1 |
| pWC0053 | pEF1a-MS2-P65-HSF1 | expressing synergistic activation mediator (SAM) | 3 |
| pWC0054 | pU6-gRNA10-MS2H | expressing gRNA10 with MS2 hairpins | 3 |
| pWC0055 | pU6-gRNA10-U6-gRNA10 | dual gRNA transcriptional units | 3 |
| pWC0056 | pCMV-2xNLS-dCas9-VPR | additional 2x nuclear localizing sequence (NLS) in the crisprTF | 3 |
| pWC0057 | p4xBS (sgRNA4)-mini-SI-mKate | encoding the synthetic operator with 4 of gRNA4 BS to drive mKate expression with SI | 3 |
| pWC0058 | p8xBS (sgRNA4)-mini-SI-mKate | encoding the synthetic operator with 8 of gRNA4 BS to drive mKate expression with SI | 3 |
| pWC0059 | p12xBS (sgRNA4)-mini-SI-mKate | encoding the synthetic operator with 12 of gRNA4 BS to drive mKate expression with SI | 3 |
| pWC0060 | p16xBS (sgRNA4)-mini-SI-mKate | encoding the synthetic operator with 16 of gRNA4 BS to drive mKate expression with SI | 3 |
| pWC0061 | p4xBS (sgRNA7)-mini-SI-mKate | encoding the synthetic operator with 4 of gRNA7 BS to drive mKate expression with SI | 3 |
| pWC0062 | p8xBS (sgRNA7)-mini-SI-mKate | encoding the synthetic operator with 8 of gRNA7 BS to drive mKate expression with SI | 3 |
| pWC0063 | p12xBS (sgRNA7)-mini-SI-mKate | encoding the synthetic operator with 12 of gRNA7 BS to drive mKate expression with SI | 3 |
| pWC0064 | p16xBS (sgRNA7)-mini-SI-mKate | encoding the synthetic operator with 16 of gRNA7 BS to drive mKate expression with SI | 3 |
| pWC0065 | p4xBS (sgRNA10)-mini-SI-mKate | encoding the synthetic operator with 4 of gRNA10 BS to drive mKate expression with SI | 3 |
| pWC0066 | p8xBS (sgRNA10)-mini-SI-mKate | encoding the synthetic operator with 8 of gRNA10 BS to drive mKate expression with SI | 3 |
| pWC0067 | p12xBS (sgRNA10)-mini-SI-mKate | encoding the synthetic operator with 12 of gRNA10 BS to drive mKate expression with SI | 3 |
| pWC0068 | p16xBS (sgRNA10)-mini-SI-mKate | encoding the synthetic operator with 16 of gRNA10 BS to drive mKate expression with SI | 3 |
| pWC0070 | pattB-puro-U6-gRNA4-2xBS-mini-mKate-CMV-dCas9-VPR | payload for BxB1-mediated genomic integration | 4 |
| pWC0071 | pattB-puro-U6-gRNA4-4xBS-mini-mKate-CMV-dCas9-VPR | payload for BxB1-mediated genomic integration | 4 |
| pWC0072 | pattB-puro-U6-gRNA4-6xBS-mini-mKate-CMV-dCas9-VPR | payload for BxB1-mediated genomic integration | 4 |
| pWC0073 | pattB-puro-U6-gRNA4-8xBS-mini-mKate-CMV-dCas9-VPR | payload for BxB1-mediated genomic integration | 4 |
| pWC0074 | pattB-puro-U6-gRNA4-12xBS-mini-mKate-CMV-dCas9-VPR | payload for BxB1-mediated genomic integration | 4 |
| pWC0075 | pattB-puro-U6-gRNA4-16xBS-mini-mKate-CMV-dCas9-VPR | payload for BxB1-mediated genomic integration | 4 |
| pWC0076 | pattB-puro-U6-gRNA4-8xBS-mini-SI-mKate-CMV-dCas9-VPR | payload for BxB1-mediated genomic integration | 4 |

| | | | |
|---|---|---|---|
| pWC0077 | pattB-puro-U6-gRNA4-16xBS-mini-SI-mKate-CMV-dCas9-VPR | payload for BxB1-mediated genomic integration | 4 |
| pWC0078 | pattB-puro-U6-gRNA10-2xBS-mini-mKate-CMV-dCas9-VPR | payload for BxB1-mediated genomic integration | 4 |
| pWC0079 | pattB-puro-U6-gRNA10-4xBS-mini-mKate-CMV-dCas9-VPR | payload for BxB1-mediated genomic integration | 4 |
| pWC0080 | pattB-puro-U6-gRNA10-6xBS-mini-mKate-CMV-dCas9-VPR | payload for BxB1-mediated genomic integration | 4 |
| pWC0081 | pattB-puro-U6-gRNA10-8xBS-mini-mKate-CMV-dCas9-VPR | payload for BxB1-mediated genomic integration | 4 |
| pWC0082 | pattB-puro-U6-gRNA10-12xBS-mini-mKate-CMV-dCas9-VPR | payload for BxB1-mediated genomic integration | 4 |
| pWC0083 | pattB-puro-U6-gRNA10-16xBS-mini-mKate-CMV-dCas9-VPR | payload for BxB1-mediated genomic integration | 4 |
| pWC0084 | pattB-puro-U6-gRNA10-8xBS-mini-SI-mKate-CMV-dCas9-VPR | payload for BxB1-mediated genomic integration | 4 |
| pWC0085 | pattB-puro-U6-gRNA10-16xBS-mini-SI-mKate-CMV-dCas9-VPR | payload for BxB1-mediated genomic integration | 4 |
| pWC0086 | pattB-puro-U6-gRNA4-8xBS-mini-mKate-CMV-dCas9-VPR-2A-bla | payload for BxB1-mediated genomic integration | 4 |
| pWC0087 | pattB-puro-U6-gRNA4-16xBS-mini-mKate-CMV-dCas9-VPR-2A-bla | payload for BxB1-mediated genomic integration | 4 |
| pWC0088 | pattB-puro-U6-gRNA4-8xBS-mini-SI-mKate-CMV-dCas9-VPR-2A-bla | payload for BxB1-mediated genomic integration | 4 |
| pWC0089 | pattB-puro-U6-gRNA4-16xBS-mini-SI-mKate-CMV-dCas9-VPR-2A-bla | payload for BxB1-mediated genomic integration | 4 |
| pWC0090 | pattB-puro-U6-gRNA10-8xBS-mini-mKate-CMV-dCas9-VPR-2A-bla | payload for BxB1-mediated genomic integration | 4 |
| pWC0091 | pattB-puro-U6-gRNA10-16xBS-mini-mKate-CMV-dCas9-VPR-2A-bla | payload for BxB1-mediated genomic integration | 4 |
| pWC0092 | pattB-puro-U6-gRNA10-8xBS-mini-SI-mKate-CMV-dCas9-VPR-2A-bla | payload for BxB1-mediated genomic integration | 4 |
| pWC0093 | pattB-puro-U6-gRNA10-16xBS-mini-SI-mKate-CMV-dCas9-VPR-2A-bla | payload for BxB1-mediated genomic integration | 4 |
| pWC0100 | pattB-hygro-CMV-dCas9-VPR-2A-G418 | dLP1-1 payload constitutively expressing dCas9-VPR | 5 |
| pWC0200 | pattB-puro-U6-gRNA10-8xBS-mini-mKate-8xBS-mini-BFP-EF1a-dCas9-VPR-2A-bla | dLP1-2 payload control expressing two reporters | 5 |
| pWC0201 | pattB-puro-U6-gRNA10-16xBS-mini-SI-mKate-16xBS-mini-SI-BFP-EF1a-dCas9-VPR-2A-bla | dLP1-2 payload control expressing two reporters | 5 |
| pWC0202 | pattB-puro-U6-gRNA10-8xBS-mini-JUG-LC-8xBS-mini-JUG-HC-EF1a-dCas9-VPR-2A-bla | dLP1-2 payload for JUG444 production (JUGAb1) | 5 |
| pWC0203 | pattB-puro-U6-gRNA10-8xBS-mini-SI-JUG-LC-8xBS-mini-SI-JUG-HC-EF1a-dCas9-VPR-2A-bla | dLP1-2 payload for JUG444 production (JUGAb2) | 5 |
| pWC0204 | pattB-puro-U6-gRNA10-16xBS-mini-JUG-LC-16xBS-mini-JUG-HC-EF1a-dCas9-VPR-2A-bla | dLP1-2 payload for JUG444 production (JUGAb3) | 5 |
| pWC0205 | pattB-puro-U6-gRNA10-16xBS-mini-SI-JUG-LC-16xBS-mini-SI-JUG-HC-EF1a-dCas9-VPR-2A-bla | dLP1-2 payload for JUG444 production (JUGAb4) | 5 |
| pWC0206 | pattB-puro-U6-gRNA10-8xBS-mini-PD1-LC-8xBS-mini-PD1-HC-EF1a-dCas9-VPR-2A-bla | dLP1-2 payload for anti-hPD1 production (PD1Ab1) | 6 |
| pWC0207 | pattB-puro-U6-gRNA10-8xBS-mini-SI-PD1-LC-8xBS-mini-SI-PD1-HC-EF1a-dCas9-VPR-2A-bla | dLP1-2 payload for anti-hPD1 production (PD1Ab2) | 6 |
| pWC0208 | pattB-puro-U6-gRNA10-16xBS-mini-PD1-LC-16xBS-mini-PD1-HC-EF1a-dCas9-VPR-2A-bla | dLP1-2 payload for anti-hPD1 production (PD1Ab3) | 6 |
| pWC0209 | pattB-puro-U6-gRNA10-16xBS-mini-SI-PD1-LC-16xBS-mini-SI-PD1-HC-EF1a-dCas9-VPR-2A-bla | dLP1-2 payload for anti-hPD1 production (PD1Ab4) | 6 |
| pWC0900 | pCMV-Gal4-VPR | constitutively expressing Gal4-VPR | S6 |
| pWC0901 | p2xUAS-BS-mini-mKate | encoding the synthetic operator with 2 of UAS BS to drive mKate expression | S6 |
| pWC0902 | p4xUAS-BS-mini-mKate | encoding the synthetic operator with 4 of UAS BS to drive mKate expression | S6 |
| pWC0903 | p6xUAS-BS-mini-mKate | encoding the synthetic operator with 6 of UAS BS to drive mKate expression | S6 |
| pWC0904 | p8xUAS-BS-mini-mKate | encoding the synthetic operator with 8 of UAS BS to drive mKate expression | S6 |
